# Pavlovian Control of Intraspinal Microstimulation to Produce Over-Ground Walking

**DOI:** 10.1101/785741

**Authors:** Ashley N Dalrymple, David A Roszko, Richard S Sutton, Vivian K Mushahwar

## Abstract

**Objective:** Neural interface technologies are more commonly used in people with neural injury. To achieve a symbiotic relationship between device and user, the control system of the device must augment remaining function and adapt to day-to-day changes. The goal of this study was to develop predictive control strategies to produce alternating, over-ground walking in a cat model of hemisection spinal cord injury (SCI) using intraspinal microstimulation (ISMS).

**Approach:** Eight cats were anaesthetized and placed in a sling over a walkway. The residual function of a hemisection SCI was mimicked by manually moving one hind-limb through the walking cycle over the walkway. ISMS targeted motor networks in the lumbosacral enlargement to activate muscles in the other limb using low levels of current (< 130 µA). Four different people took turns to move the “intact” limb. Two control strategies, which used ground reaction force and angular velocity information about the manually moved limb to control the timing of the transitions of the other limb, were compared. The first strategy, reaction-based control, used thresholds on the sensor values to initiate state transitions. The other strategy used a reinforcement learning strategy, Pavlovian control, to learn predictions about the sensor values. Thresholds on the predictions were used to initiate transitions.

**Main Results:** Both control strategies were able to produce alternating, over-ground walking. Reaction-based control required manual tuning of the thresholds for each person to produce walking, whereas Pavlovian control did not. We demonstrate that learning occurs quickly during walking. Predictions of the sensor signals were learned quickly, initiating transitions in no more than 4 steps. Pavlovian control was resilient to transitions between people walking the limb, between cat experiments, and recovered from mistakes during walking.

**Significance:** This work demonstrates that Pavlovian control can augment remaining function and allow for personalized walking with minimal tuning requirements.

## INTRODUCTION

After a spinal cord injury (SCI), people experience motor and sensory paralysis to varying degrees, depending on the severity and level of the injury. Two-thirds of all SCIs in the USA are incomplete (“Spinal Cord Injury (SCI) 2017 Facts and Figures at a Glance” 2017). For people with paraplegia, regaining the ability to walk is a high priority, ranking first or second nearly 40% of the time (Anderson 2004). Currently, SCI has no cure; therefore, regaining the ability to walk has been pursued through other means such as rehabilitation (Musselman et al. 2009; Lam et al. 2015; Morrison et al. 2018), neural technologies (Kobetic et al., 1997; Hardin et al. 2007; Moritz et al., 2008; Holinski et al. 2016), or a combinatorial approach (Carhart et al. 2004; Angeli et al. 2018; Gill et al. 2018).

The neural networks in the spinal cord below the SCI and their connections to the leg muscles remain intact (Hunter and Ashby 1994). These spinal networks can be targeted and activated using electrical stimulation (Mushahwar and Horch 2000; Saigal et al., 2004; Hofstoetter et al. 2015; Angeli et al. 2018; Wagner et al. 2018; Gill et al. 2018). One type of electrical stimulation technique is intraspinal microstimulation (ISMS), which entails implanting fine, hair-like microwires in the ventral horn of the lumbosacral enlargement. Interestingly, stimulation in this region through a single microwire produces large graded single joint movements as well as coordinated multi-joint synergies (Mushahwar and Horch 2000; Saigal et al., 2004; Mushahwar and Horch 1998; Holinski et al. 2011). Through targeted activation of hind-limb muscles, ISMS has been used to restore walking in anaesthetized (Holinski et al. 2016; 2013) and spinalized cats (Saigal et al., 2004). Recently, ISMS was used in cats to produce nearly 1 km of over-ground, weight-bearing walking (Holinski et al. 2016). These distances were achieved immediately after implantation of the microdevice and without the need for extensive rehabilitation. Responses produced by ISMS remain consistent throughout the use of the implant (Mushahwar et al., 2000), and long-term use of ISMS for walking will likely further improve walking distances achieved. Therefore, ISMS is poised to be a viable clinical approach to restoring walking after severe paralysis. An important and clinically-relevant aspect of a successful neural prosthesis is the control of the device and how users interact with the control strategy.

Current commercially available devices for restoring walking after SCI, such as the Parastep, Praxis, and various exoskeletons, have limited control options. The Parastep and Praxis systems use surface and implanted functional electrical stimulation (FES) electrodes, respectively (Chaplin 1996; Johnston et al. 2005). Walking is accomplished using open loop alternation between stimulation of the quadriceps muscles and the peroneal nerve, with each step initiated using push-buttons on a walker. Powered exoskeletons initiate open-loop walking by the user leaning forward (Chang et al. 2015; Ekelem and Goldfarb 2018). The users are expected to adapt their walking to accommodate the control strategy in the device. To restore meaningful and functional walking, especially after an incomplete SCI, the control strategy needs to adapt to the user, utilize residual function, and deliver stimulation to compensate for the deficits as needed.

Controllers developed for ISMS to date have primarily focused on restoring walking in models of complete SCI (Dalrymple and Mushahwar 2017). Feedback, such as ground reaction force, hip angle, or activity of sensory neurons from the dorsal root ganglia, were used to modify the inherent timing of the transitions between the phases of the gait cycle (Saigal et al. 2004; Holinski et al. 2011; Holinski et al. 2013, 2016). A recent paper depicted the first control strategies developed for ISMS in a model of incomplete SCI (Dalrymple et al. 2018). These strategies augmented the residual function in a model of hemisection SCI and, using supervised machine learning, adapted the control strategy for different speeds of walking. However, as people with SCI experience varying levels of paralysis, each person would require their own custom stimulation settings to restore walking. Moreover, incomplete injuries evolve over time requiring further updating of stimulation settings. Manual tuning of settings is burdensome; it is time-consuming and based qualitatively on trial and error.

Control strategies utilizing machine learning are needed for automatic adaptation of stimulation settings to restore walking. Supervised machine learning has been used to control surface functional electrical stimulation (FES) systems in persons with SCI to track joint angles (Abbas and Triolo 1997; Popović et al. 1999; Qi et al. 1999), initiate the swing phase (Kirkwood and Andrews 1989; Kostov et al. 1992, 1995; Tong and Granat 1999; Sepulveda et al. 1997), control FES over multiple joints (Fisekovic and Popovic 2001), predict different phases of the gait cycle in able-bodied subjects (Kirkwood and Andrews 1989; Williamson and Andrews 2000), and in finite control of FES walking after complete SCI (Popović 1993). However, supervised learning requires manual labelling of data and is limited by the data set used for training. Many examples with sufficient variability are needed in the training data set to obtain an accurate generalization. Ideally, stimulation settings would be tuned once during the initial set-up for each person, and thereafter automatically adjust to any changes in daily gait patterns. A recent machine learning approach demonstrated the feasibility of adaptive tuning of impedance parameters in a prosthetic knee (Wen et al. 2019); however, to date, machine learning approaches have not been utilized in implanted neural prosthetic approaches for restoring over-ground walking.

Intuitive control of a neural prosthesis requires the device to know what the user wants to do pre-emptively with automatic adaptation to changes in the environment. Learning predictions of walking-relevant sensor signals for initiating control outputs may be a more reliable method to produce walking. Predictions allow for a timely response that can be modified with experience. In this study, we compared more traditional control methods with a new prediction-based machine learning control method, called Pavlovian control, to produce over-ground, alternating walking in a model of hemisection SCI. Specifically, we assessed the need for manual tuning of control settings between reaction-based control and Pavlovian control over several cat experiments and with different people participating to move one limb through the walking cycle and after perturbations. This presents the first application of Pavlovian control to produce walking. It is also the first known application of RL techniques in a spinal neural interface. Using Pavlovian control, we demonstrate that alternating over-ground walking can be achieved quickly using predictions of walking-relevant sensor signals, and that the thresholds for Pavlovian control do not require re-tuning across different conditions.

## METHODS

All experimental procedures were approved by the University of Alberta Animal Care and Use Committee under protocol AUP301. Eight adult male cats (3.96 to 5.22 kg) were individually housed in large cages and were provided with daily enrichment that included a larger play pen, toys, human interaction, and soothing music.

### Implant Procedure

Investigations were conducted in acute, non-recovery experiments. Anaesthesia was initially induced with isoflurane (5%), while all surgical procedures and data collections were performed under sodium pentobarbital anesthesia administered intravenously (induction: 25mg/kg; maintenance: 1 in 10 dilutions in saline). A laminectomy was performed to expose the lumbosacral enlargement. An array of 12 microwire electrodes made of Pt-Ir (80/20), 50 µm in diameter, insulated with 4 µm polyimide except for approximately 400 µm exposure at the tip, was implanted in one side of the spinal cord according to established procedures (Mushahwar et al. 2000; Bamford et al. 2016). The microwire tips targeted lamina IX in the ventral horn based on functional maps of the motoneuron pools (Mushahwar and Horch 2000; 1998; Vanderhorst and Holstege 1997). In addition to motoneuronal pools, this region contains neural networks that, when stimulated, produce coordinated multi-joint synergistic movements of the leg (Holinski et al. 2016; Mushahwar and Horch 2000; Saigal et al., 2004; Bhumbra and Beato 2018; Engberg and Lundberg 1969).

### Stimulation Protocol

Trains of stimuli were delivered using a customized current-controlled stimulator (Sigenics Inc., Chicago, IL, USA) and consisted of a trapezoidal waveform that ramped from threshold to chosen amplitude over 3 time-steps (time-step = 40 ms). The stimulus pulses in the trains were 290 µs in duration, biphasic, charge-balanced and delivered at a rate of 50 Hz. Stimulation amplitudes ranged from threshold (< 20 µA) to amplitudes that produced weight-bearing movements (60 to 80 µA) and did not exceed 130 µA through any electrode.

The movements elicited by stimulation through single electrodes were hip flexion, hip extension, knee extension, ankle dorsiflexion, ankle plantarflexion, and a backward extensor synergy, which were combined to construct a full walking cycle. Of the 12 electrodes implanted unilaterally, between 5 and 9 were needed to produce the desired walking movements and included redundancy in the functional targets. Stimulation channels were combined to construct the four phases of the walking cycle: F (early swing), E1 (late swing to paw-touch), E2 (mid-stance), and E3 (propulsion) (Goslow et al. 1973; Engberg and Lundberg 1969). The phases F - E2 and E1 - E3 were defined to be opposite phases of the walking cycle.

### Experimental Setup

Following the implantation of the ISMS array, cats were transferred to a custom-built instrumented walkway and placed in a sling that supported their trunk, head and forelimbs. The hind-limbs where left to move freely over the walkway (Figure 1). The cats remained anesthetized for the duration of the experiment. The sling was fixed on a cart that moved with the cat over the walkway. The cart was partially unloaded to offset the weight of the recording and stimulating equipment as well as mobile vital signs monitors that were placed on it.

**Figure 1.**
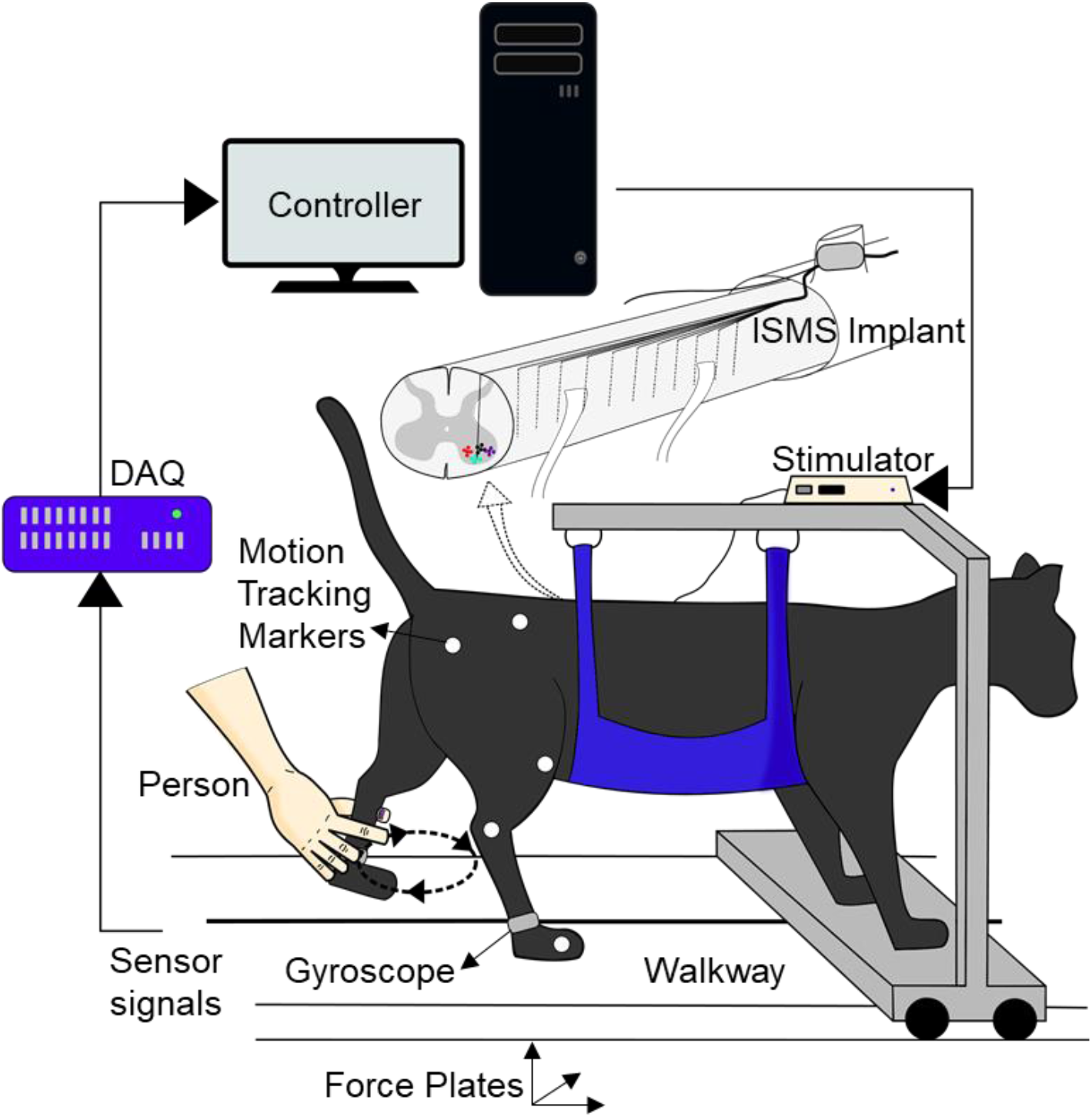
Experimental setup for over-ground walking. A naïve experimenter moved the left hind-limb through the walking cycle. Sensor signals from force plates under the walkway and a gyroscope on the tarsals from both hind-limbs were converted to digital signals by the DAQ (data acquisition device) and streamed into Matlab. In Matlab, a custom control algorithm was used to control the stimulation to the spinal cord to move the right hind-limb to the opposite phase of the walking cycle.

Gyroscopes were placed on the tarsals of each hind-limb to measure angular velocity in real time. Three-dimensional force plates were mounted underneath the walkway and used to measure vertical ground reaction forces of each limb. The sensor signals were filtered using a Butterworth filter (f_C_ = 3 Hz, 2^nd^ order) and digitized at 1 kHz using the Grapevine Neural Interface Processor (Ripple, Salt Lake City, UT, USA) and streamed into Matlab (MathWorks, Inc., Natick, MA, USA) during walking.

Reflective markers were positioned on the iliac crest, hip, knee, ankle, and metatarsophalangeal (MTP) joints of the right hind-limb. Kinematics of this limb were recorded using a camera (120fps, JVC Americas Corp., Wayne, NJ, USA) positioned 4.5 m away from the center of the walkway. Marker positions were tracked post-hoc using MotionTracker2D, a custom Matlab program written by Dr. Douglas Weber (University of Pittsburgh, Pittsburgh, PA, USA).

A hemisection SCI was modeled in anaesthetized cats with an intact spinal cord. A person (naïve experimenter) manually moved the left hind-limb through the walking cycle (person-moved limb; PML) to represent the intact leg, while the right hind-limb was moved using ISMS (stimulation-controlled limb; SCL) and represented the paralyzed leg (Figure 1). This hemisection SCI model is similar to Brown-Sequard syndrome in humans, where one leg is paralyzed and the other is motor-intact (Kunam et al. 2018).

### Control Strategies

The goal of the control strategies was to transition the SCL through the walking cycle such that the phase of the SCL was opposite to the phase of the PML and that the force produced by the SCL was enough to propel the animal across the walkway to produce over-ground walking. Each control strategy determined when the SCL transitioned from one phase to the next based on sensor information from the PML.

#### Reaction-based Control Strategy

For reaction-based control, thresholds were placed on the sensor signals recorded from the PML during walking to trigger transitions between the phases of the walking cycle in the SCL. The sensor signals used for defining the transitions between phases of the walking cycle were ground reaction force and angular velocity of the PML (Figure 2A). The transitions were controlled by rules involving the current phase in the walking cycle, comparing the sensor values with threshold values, and the direction of the slope of the sensor values. Thresholds were placed on the sensor signals such that they anticipated when the transitions would normally occur to account for the electromechanical delay of approximately 200 ms (Dalrymple et al. 2018).

**Figure 2.**
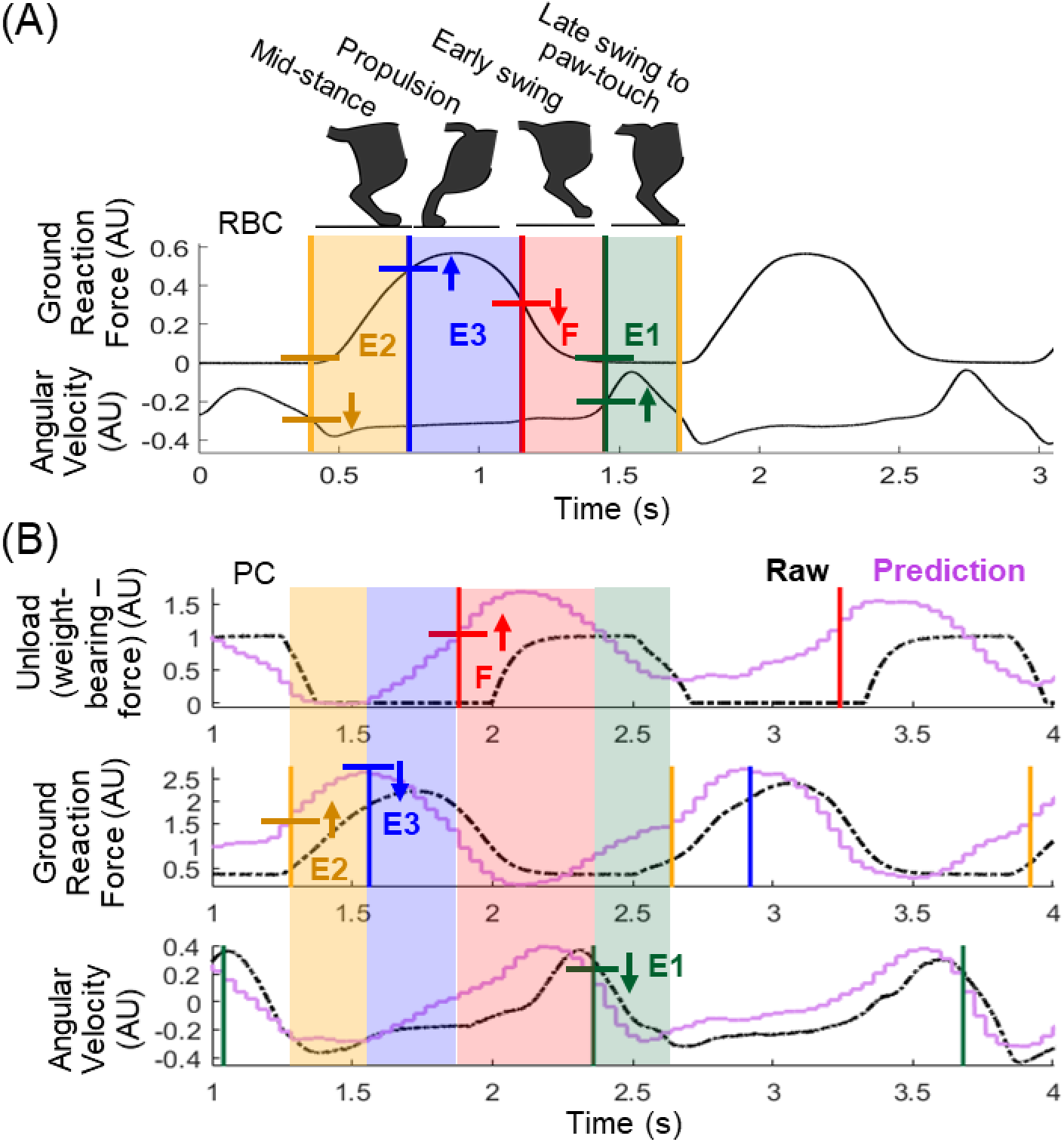
**(A)** Phases of the walking cycle aligned with phases (E2, E3, F, E1) and thresholds on predictions of sensor signals from the PML (person-moved limb) for rule-based control (RBS). (B) Threshold settings for one naïve experimenter (person A) on raw data from the PML using Pavlovian control (PC). AU = arbitrary units due to normalization during acquisition. Shaded regions indicate the phase of the walking cycle detected on the PML, divided by vertical lines indicating timing of transitions. Horizontal lines mark the threshold values for corresponding phase. Arrows indicate the direction of the slope of the signal required by the algorithm.

#### Pavlovian Control Strategy

Automatic adaptation of the control output is required to produce personalized walking that augments remaining function after a SCI. We used novel reinforcement learning methods to predict three walking-relevant signals in real time: the ground reaction force, angular velocity, and unloading.

Pavlovian control borrows concepts from Pavlovian conditioning to use learned predictions to trigger fixed or pre-defined outputs, such as a stimulation output (Modayil and Sutton 2014). When applied to a control problem, first, predictions of sensory stimuli must be learned. Then the learned predictions trigger a fixed output response. Reinforcement learning (RL) was used to learn the predictions of three walking-relevant signals of the PML. The signals of interest were the ground reaction force, angular velocity, and unloading of the PML. Unloading was defined as the weight-bearing threshold (equal to 12.5% of the cat’s body weight in this setup (Lau et al. 2007)) minus the ground reaction force. Unloading differs from ground reaction force as it informs when the PML is below or above a weight-bearing threshold. Thresholds were placed on the predictions of these signals, which in addition to the slope of the predictions and knowledge of the current phase were used to transition the SCL to the opposite phase of the PML (Figure 2B).

#### Learning Methods

##### State Representation of Sensor Signals

Sensor signals are complex with a wide range of possible values. Learning requires a combination of sensor values to be repeated multiple times. With highly sampled, broad ranges of values for multiple sensors, exact duplicates of overlapping sensor values are unlikely to occur, making learning slow. Therefore, it is necessary to generalize the state space of sensor signals by function approximation. This process converts the complex, high-dimensional sensor data into a binary vector representation of the state space, named the feature vector, **x**.

Six sensors were chosen to form the state space: left ground reaction force, right ground reaction force, the sum of left and right ground reaction forces, left angular velocity, right angular velocity, and the exponential moving average of the left ground reaction force. The exponential moving average gives a long-term history of the force signal and helps differentiate between the periodic increasing and decreasing of the other sensor signals. First, the sensor values were normalized from their usable range to values between 0 and 1. Selective Kanerva coding was used to represent the normalized sensor values as a binary vector (Travnik and Pilarski 2017).

To perform selective Kanerva coding, *K* = 5000 specific states, also referred to as prototypes, were randomly distributed over the entire normalized, 6-dimensional state space (6 sensors; Figure 3). The prototype locations were held constant for all experiments. Hoare’s quickselect was used to find the *c* closest prototypes to the current state according to their Euclidean distance. Three values of *c*, determined by choosing small ratios, η, such that *c = K*η were used. These values of *c* equaled to 500, 125, and 25, corresponding to η values of 0.1, 0.025, and 0.005, respectively. Using multiple *c* values is similar to the use of overlapping tilings in tile coding (Sutton and Barto 2018); it allows for coarse and fine representation of the state in the feature vector. When a combination of sensors values occurred, defining the current state in the state space, the *c*-closest features were activated in the feature vector (set equal to 1), while the rest equal 0. The total number of features in **x** was 3*K*, where 650 (*c_1_ + c_2_ + c_3_*) features were active at all times. The pseudocode for selective Kanerva coding used in this work is provided in Algorithm 1. Bolded variables refer to vectors or matrices; italicized variables refer to constants with values that pertain to this work.

**Figure 3.**
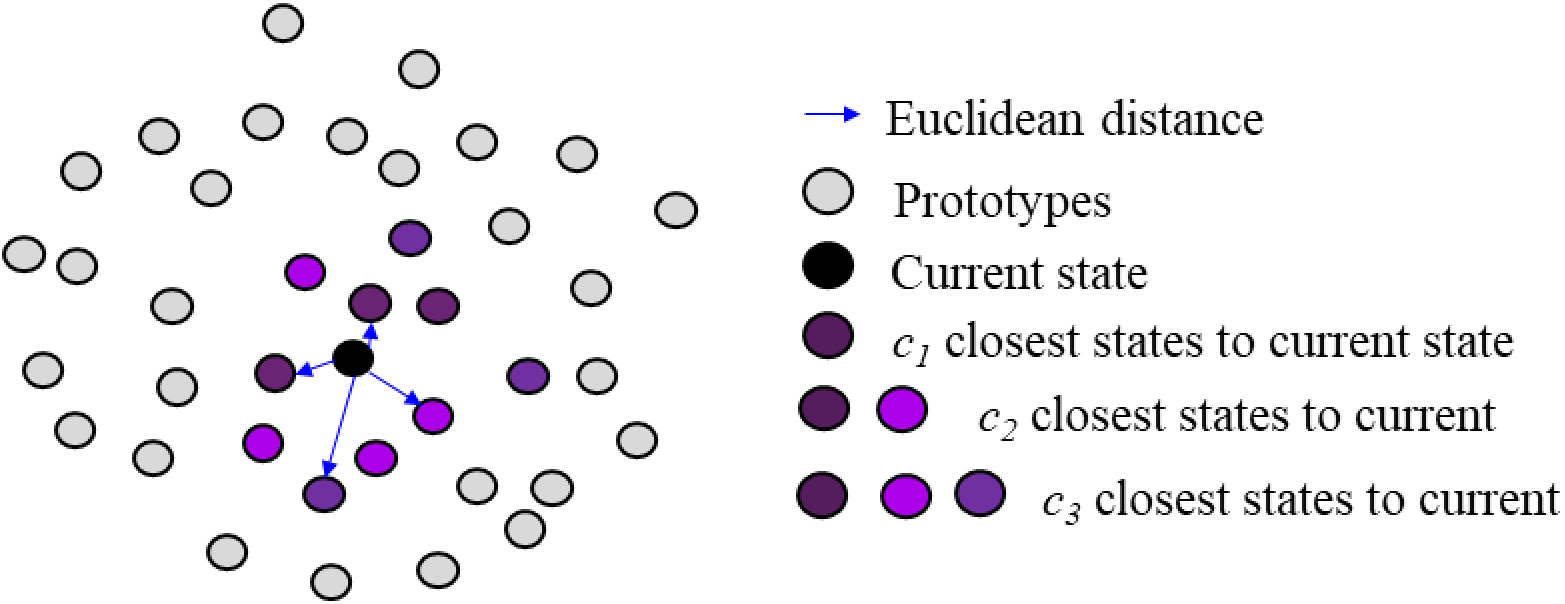
A depiction of selective Kanerva coding (SKC). Prototypes closest to the current state within the state space are activated.

**Algorithm 1.**
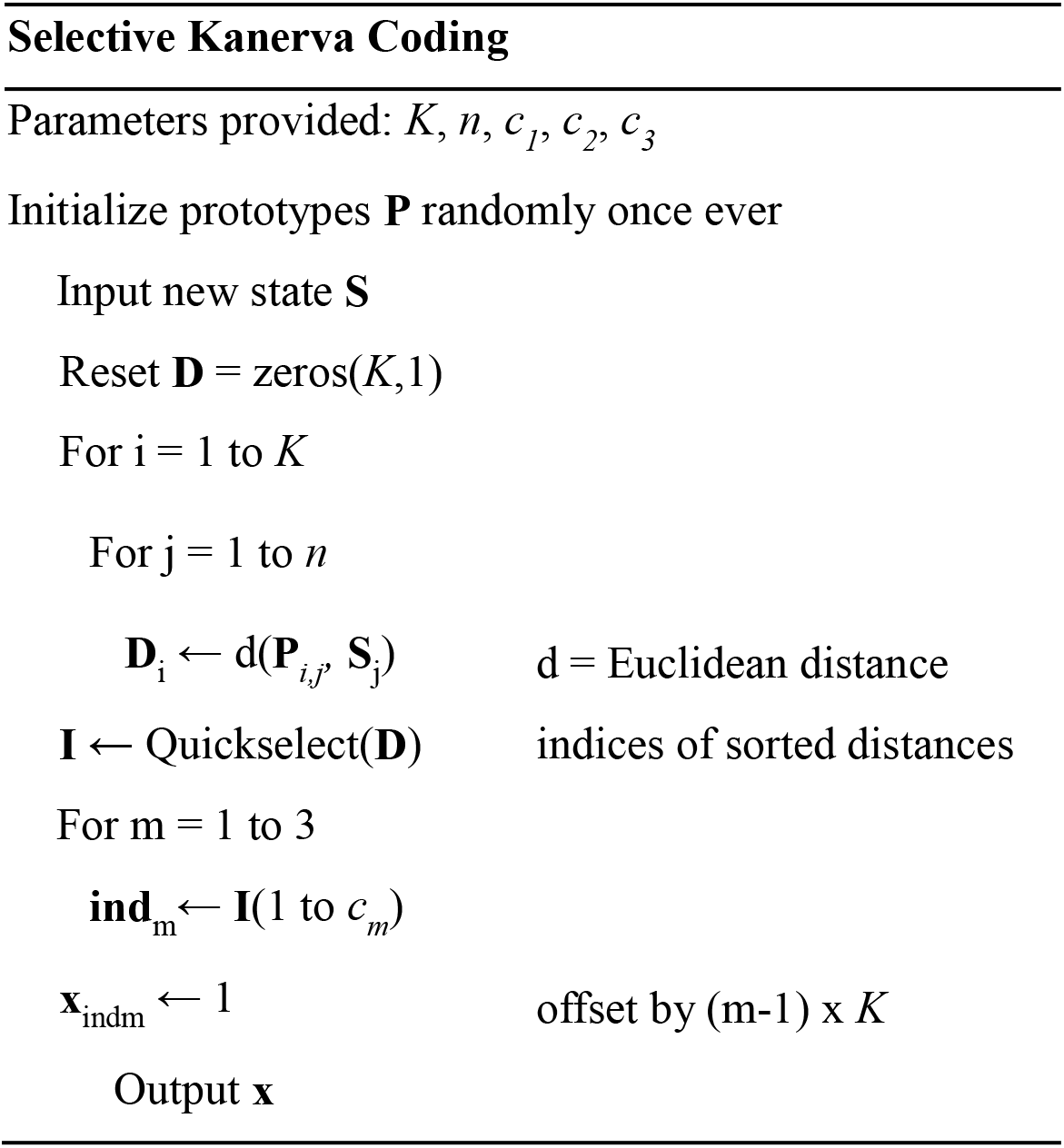
Selective Kanerva Coding as used in this work. Function approximation method to convert the sensor state space into a binary feature vector for use in reinforcement learning.

##### True Online Temporal Difference Learning

True online temporal difference learning (TOTD), which is a reinforcement learning method, was used to learn the predictions of the sensor signals in real time during walking. Similar to operant conditioning, RL is an area of machine learning that accomplishes a goal by maximizing future reward (Skinner 1963; Staddon and Cerutti 2003; Sutton and Barto 2018). RL can also estimate, or predict, the future values of signals other than reward. General value functions (GVFs) can be learned to predict arbitrary signals of interest, called cumulants (Z) (White 2015). Many GVFs can be learned simultaneously to produce predictions of many cumulants. Temporal difference (TD) learning is a method that can be used to estimate the future values of cumulants using previously obtained estimates, called bootstrapping (Sutton 1988; van Seijen et al. 2015; Sutton and Barto 2018). TOTD is a newer method that matches the forward view of temporal difference learning online exactly by adding terms to the eligibility trace and weight update equations (van Seijen and Sutton 2014; van Seijen et al. 2015).

During walking, TOTD predicted the future values of three signals recorded from the PML: unloading, ground reaction force, and angular velocity. Specifically, the returns of the cumulants were estimated in real time through the inner product of the weight vector (updated during TOTD) and the feature vector from function approximation (selective Kanerva coding), to produce the GVF for that cumulant (Algorithm 2). The learning step-size (α), which determines the magnitude of the update, was set to 0.001, which was determined empirically. The bootstrapping parameter for the eligibility trace (λ) was set to 0.9 as is often standard. Different termination signals (γ) were determined for each cumulant empirically: 0.9 for unloading, 0.71 for ground reaction force, and 0.75 for the angular velocity. As 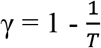, where *T* = 40 ms (one time-step), these values corresponded to timescales of 400 ms, 138 ms, and 160 ms, respectively.

**Algorithm 2.**
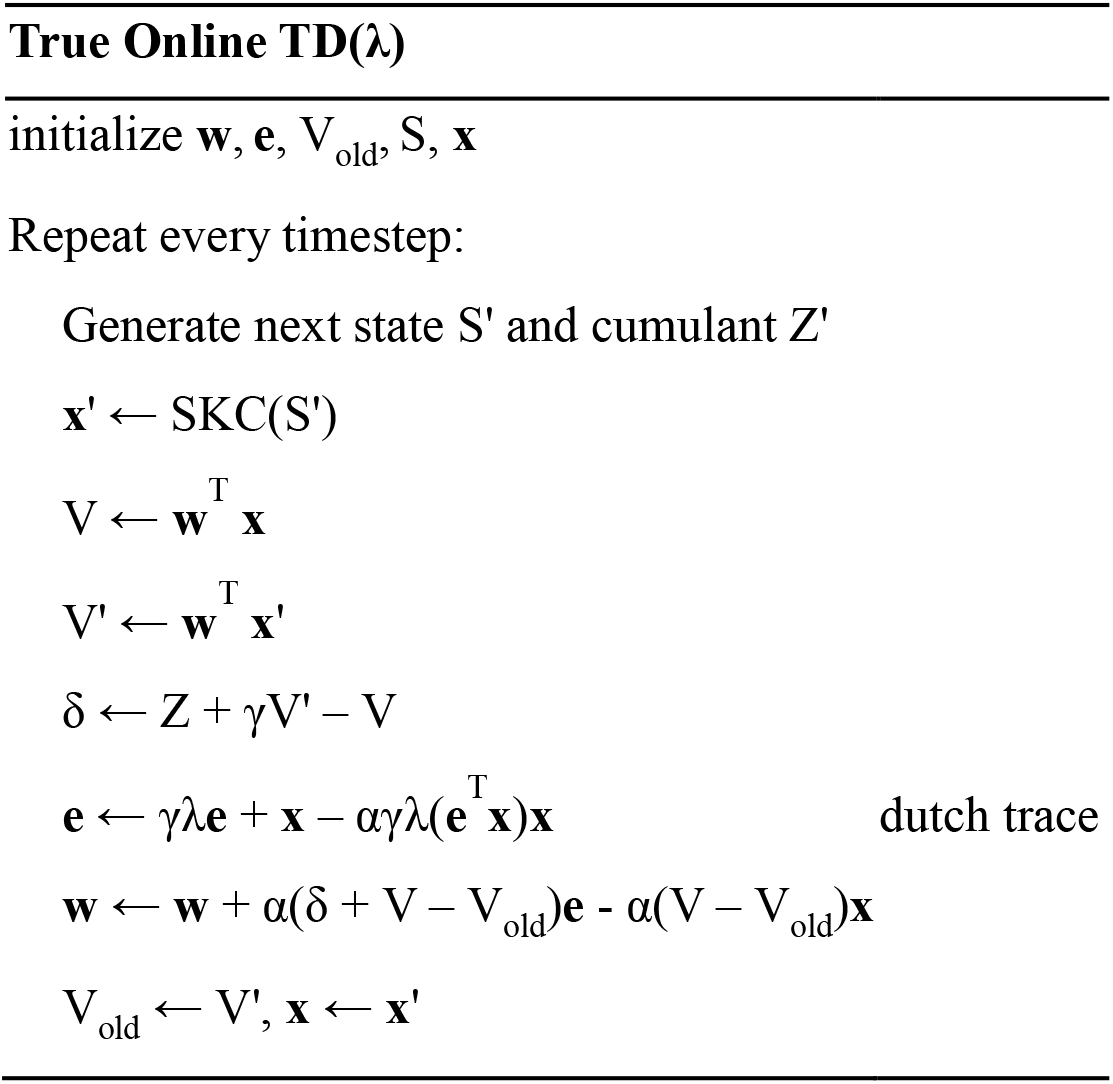
True online temporal difference learning. Reinforcement learning algorithm to estimate the discounted future values of sensor signals during walking.

Thresholds were placed on the GVFs, which in addition to the direction of the slope of the GVF and the current phase of the walking cycle of the PML, triggered transitions between the phases of the walking cycle of the SCL to be in the opposite state of the PML (Figure 2B). The prediction of a sensor value produced a fixed stimulation response (the stimulation parameters did not vary during walking), thus utilizing Pavlovian control to produce over-ground walking. The phase transitions were triggered by the raw sensor values crossing a threshold (unconditioned stimulus) if the predicted value (conditioned stimulus) did not elicit a response. These are referred to as back-up reactions. The thresholds for the back-up reactions were held constant throughout all walking trials.

Of the 8 cat experiments conducted in this study, the first 3 had one set of thresholds on the GVFs, while the remaining 5 had a different set of thresholds. The learning parameters and methods remained constant throughout the study. The initial thresholds for Pavlovian control were chosen based on testing on previously collected data from treadmill stepping (Dalrymple et al. 2018) and bench testing on the walkway without a cat. These thresholds resulted in 55.4% of the steps triggered by back-up reactions and 2.0% of steps having missed phase transitions. Therefore, the thresholds were revised along with a change in which signal was used to predict some of the phases and held constant for the following 5 experiments. The back-up reaction thresholds were unchanged.

### Experimental Protocol

A walking trial consisted of one trip across the walkway (∼ 3 m). A naïve experimenter manually moved the PML through the walking cycle. The SCL pushed the anaesthetized cat and cart across the walkway. Up to four different naïve experimenters moved the limb through the walking cycle in each experiment. The control method (reaction-based or Pavlovian) used for each walking trial was determined randomly by a different person than the one walking the PML, or by a random number generator. The person moving the limb was blinded to the control method driving ISMS for each trial.

For some trials, experimenters were told to purposefully make a mistake while walking the PML. A mistake was not explicitly defined; it was left to the discretion of the person walking the PML. Intentional mistakes included elongating the stance or the swing phase, shaking the limb in the air, or slipping forward or backward.

#### Reaction-based Control Trials

The phase transition thresholds for each naïve experimenter moving the PML were based on sensor values they produced during 2 consecutive walking trials. The person-specific thresholds remained constant throughout all cat experiments. Each person performed walking trials using the customized thresholds from the three other naïve experimenters walking the PML in addition to trials with their own thresholds.

#### Pavlovian Control Trials

Several different trial types were conducted to investigate early learning, continued learning, and how the learning adapted or recovered after changes between cat experiments and people walking the PML. Early learning was evaluated by initializing the learning weights, eligibility trace, and GVFs to 0 at the beginning of a walking trial. In these trials, learning began anew with no prior knowledge. These early learning trials were repeated in every cat experiment with different naïve experimenters walking the PML.

Learning also continued across several walking trials within each cat experiment. Throughout these trials within the experiment, multiple naïve experimenters took turns to walk the PML through the walking cycle. Furthermore, the carry-over of learning from one cat experiment to the next was tested over 5 cats. Repeating these carry-over trials in a new cat experiment allowed repeated investigation of the transfer of learning between experiments with different cats and experimenters walking the PML. A set of trials were also conducted whereby learning continued throughout 5 cat experiments, where multiple experimenters took turns to walk the PML within each experiment. These trials investigated the long-term learning and the adaptation to changes in cats and people walking the PML.

### Statistics

A one-sample t-test was used to compare the alternation phase differences with the target of 180°. A p-value ≤ 0.05 was considered to indicate significance. The effect size was determined using Cohen’s d.

**Χ**^2^ tests were conducted to compare the proportion of prediction-triggered phase transitions between different Pavlovian control walking trial types (early, within one cat, carry-over, and continued learning), as well as for comparing the proportion of missed phase transitions across control methods. Cross-tabulations were generated for all pair-wise combinations. The **Χ**^2^ with the continuity correction for 2 × 2 contingency tables were reported, with the α-level adjusted using the modified Bonferroni correction for multiple comparisons.

### Data Processing and Analysis

#### Calculating Alternation

The alternation of the two hind-limbs was calculated from the ground reaction forces using previously described methods (Dalrymple et al. 2018). Briefly, the time spent in loading per leg was converted into the degrees of a circle, with the onset of loading of the PML defining the points of 0° and 360°. The half-way time of loading for each limb was converted to degrees according to the step period. The difference of the phase for each limb should equal 180° for perfect alternation.

#### Defining Transitions as Triggered by a Prediction or a Reaction

A step was considered to be entirely under Pavlovian control if all 4 phases of the gait cycle were transitioned using the prediction crossing the threshold. If any of the phases required a back-up reaction to transition, then that entire step was counted as such.

#### Learning Curves

The online prediction of the return (predicted discounted future values of the sensor signals) with the ideal return (actual discounted sum of future values of the sensor signals) was compared for each sensor value (Sutton and Barto 2018). During walking, TOTD estimated the return based on previous interaction and current sensor values. The ideal return was calculated post-hoc by summating the future raw sensor values discounted by the discount factor (γ) used for each sensor signal. The mean squared error between the online return and the ideal return for each sensor signal was calculated for early learning trials and averaged the errors over the trial time.

## RESULTS

A total of 7943 steps from 770 trials were recorded from eight cats. On average, the step period was 1.32 s (SD = 0.26 s) and ranged from 0.44 s to 2.82 s.

### Walking with Reaction-Based Control

The translatability of the tuned parameters was tested from one walking pattern (by one naïve experimenter) to another and the need for retuning of parameters for reaction-based control of over-ground walking was. Reaction-based control was tested in all eight cats, resulting in 264 walking trials. There was high variability in the force production and movements produced by the 4 naïve experimenters walking the PML (Figure 4A); therefore, customized thresholds for transitions between the phases of the walking cycle for each person were needed. One naïve experimenter’s thresholds were not translatable to the other naïve experimenters. The best performance of customized thresholds was 89.6% of steps successfully transitioning through the phases of the walking cycle. Overall, 18.7% (680/3645) of the total number of steps in all walking trials under reaction-based control had missed phase transitions due to the inability of the sensor values to cross the thresholds (Table 1).

**Figure 4.**
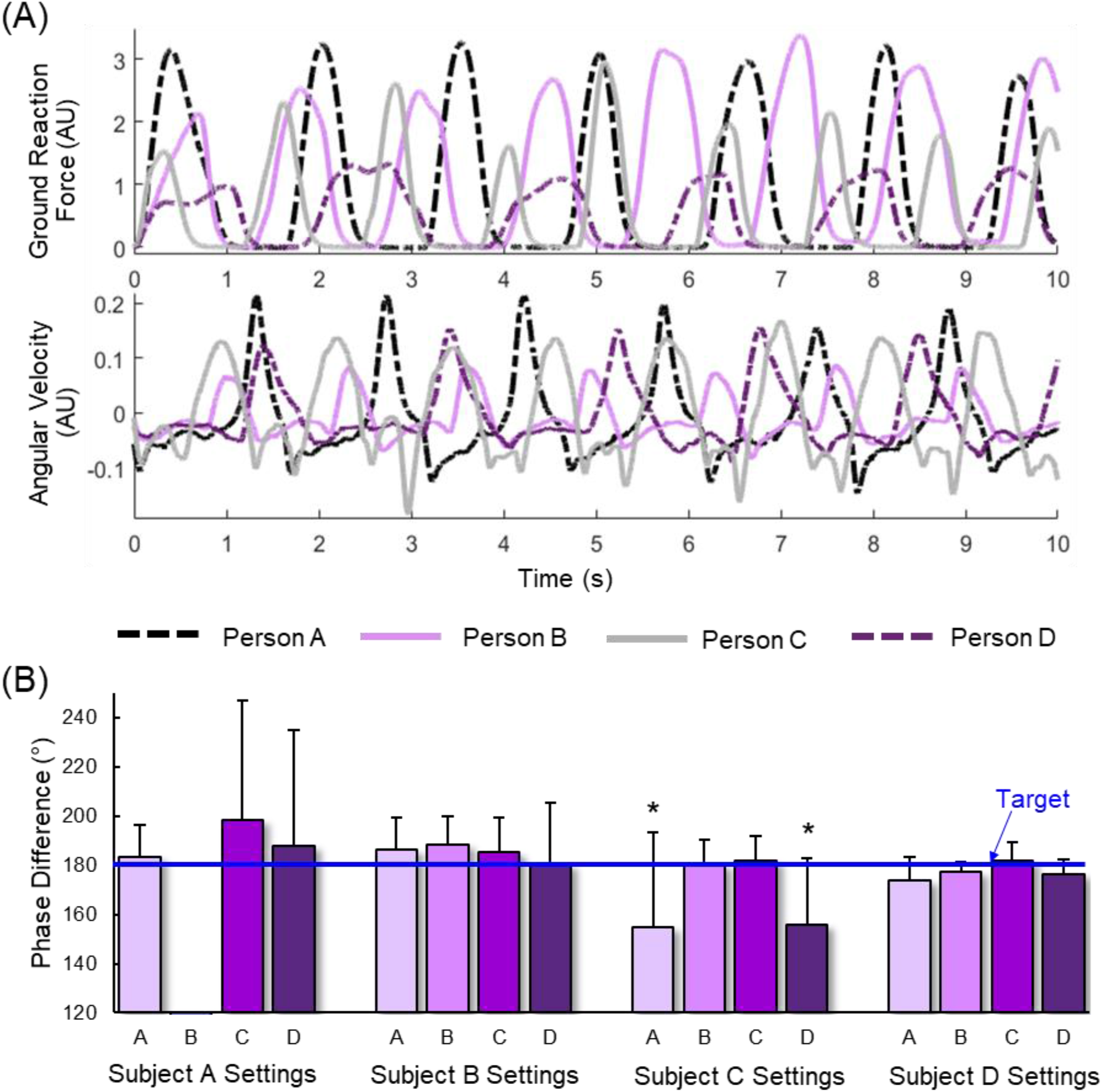
Walking using reaction-based control (n = 264 trials). **(A)** Ground reaction forces and angular velocities produced by each of the 4 naïve experimenters walking the PML (person-moved limb). **(B)** Alternation phase differences of the hind-limbs for each person walking the PML with threshold settings tuned for each person. Target alternation is 180°. * p < 0.0001.

**Table 1.** Proportion of missed steps for combinations of people walking the PML (person-moved limb) using customized threshold settings for each person during reaction-based control.

| Missed Steps using Reaction-Based Control (%) |  |  |  |  |  |
| --- | --- | --- | --- | --- | --- |
| Threshold Settings | Person Walking Limb |  |  |  |  |
|  |  | A | B | C | D |
|  | A | 11.0% | 100% | 94.0% | 54.8% |
|  | B | 7.1% | 10.4% | 46.6% | 12.5% |
|  | C | 5.9% | 44.1% | 12.0% | 6.0% |
|  | D | 12.4% | 12.8% | 42.0% | 11.5% |

The alternation between the PML and the SCL was assessed using their phase difference, where a phase difference of 180° indicated perfect alternation (Dalrymple et al. 2018). Both hind-limb alternation and successful transitions through the phases of the walking cycle must occur for walking to be considered functionally effective. Overall, reaction-based control achieved a phase difference of 179.6° (SD = 19.0°). There were instances where the parameter settings for one naïve experimenter (person C) walking the PML resulted in fewer missed phase transitions when utilized for another experimenter walking the PML (persons A and D; Table 1). However, poor PML-SCL alternation was encountered due to large variability in the walking patterns produced by the different experimenters (Figure 4B). This was because persons A and D made larger movements with larger sensor values than required for the settings tuned for person C, triggering phase transitions between the phases of the gait cycle earlier than needed to produce alternating walking. This produced a phase difference significantly less than 180° with very large effect sizes (phase difference for A = 155.0°; phase difference for D = 155.9°; p < 0.0001; df = 173, 46; one-sample t-test; Cohen’s d = 0.64, 1.97). The inconsistent alternation and unsuccessful phase transitions across different people walking the PML (i.e., different walking patterns) highlight the need for an automatically adapting control system.

### Walking with Pavlovian Control

#### Learning to predict sensor signals occurs quickly to produce over-ground walking

The Pavlovian controller learned predictions in real time during over-ground walking. Without prior learning, predictions became the only signals that initiated proper phase transitions within a maximum of 4 steps, which corresponded to approximately 4 s. Back-up reactions for phase transitions most commonly occurred within the first step compared to later steps, indicating that learning the predicted signals occurred quickly to initiate phase transitions (Table 2). Fast learning is also demonstrated by the learning curves, where the mean squared error between the online and ideal returns decreases exponentially as learning continues within the trial (Figure 5). Throughout all 1036 steps in the early learning trials, only 3 had failed phase transitions throughout the walking cycle. In 87.2% of the steps taken, the phase transitions were initiated by the predictions crossing the thresholds. Early learning trials had an average phase difference of 181.9° ± 7.8°. Therefore, the naïve learning algorithm was able to quickly learn accurate predictions of walking-relevant sensor signals to produce over-ground walking using ISMS.

**Figure 5.**
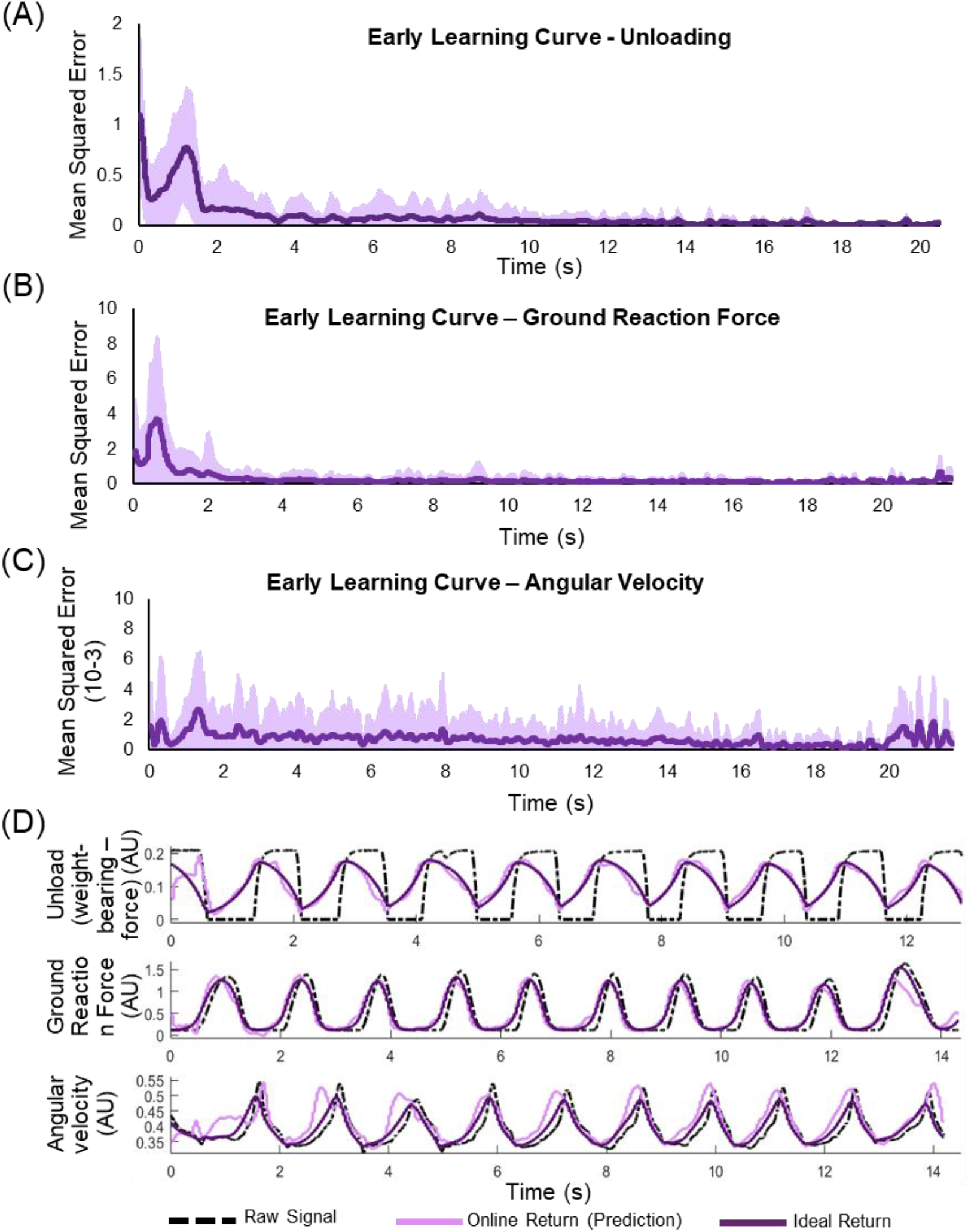
Characterizing speed of early learning (n = 88 trials). **(A)** Learning curve depicting the average (solid line) and standard deviation (shaded region) of the mean squared error of the unloading signal over time. **(B)** Learning curve for ground reaction force. **(C)** Learning curve for angular velocity. **(D)** Example of the actual sensor signals, online estimated return (prediction), and the ideal return (actual discounted sum of future values of the sensor signals).

**Table 2.** Back-up reactions in early learning trials using Pavlovian control. Within how many steps at the beginning of a walking trial was a back-up reaction triggered for each person walking the PML (person-moved limb).

| Reactions in Early Learning Trials |  |  |  |  |
| --- | --- | --- | --- | --- |
|  | 1 Step | 2 Steps | 3 Steps | More |
| A | 92.3% | 3.8% | 3.8% | 0.0% |
| B | 97.1% | 2.9% | 0.0% | 0.0% |
| C | 69.6% | 8.7% | 17.4% | 4.3% |
| D | 20.0% | 0.0% | 40.0% | 40.0% |
| All | 84.1% | 4.5% | 8.0% | 3.4% |

#### Learning that continued within a cat experiment produced better Pavlovian control

As learning continued past one walking trial within a cat experiment, the predictions of the walking-relevant signals became smoother and more reliable as they accumulated more experience (Figure 6A-B). The proportion of steps initiated by predictions crossing the thresholds significantly increased compared to trials without prior learning (initialized to zero: 87.4% predictions; continued within one cat: 95.6% predictions; p < 0.0001, **Χ**^2^ test; Figure 6D). The phase difference achieved in these trials was 181.1° and was not significantly different from the target of 180° ± 5.9° (p = 0.077; df = 98; one-sample t-test), demonstrating the ability to maintain alternation of the hind-limbs as online learning continued within a cat experiment for all experimenters walking the PML (Figure 6E).

**Figure 6.**
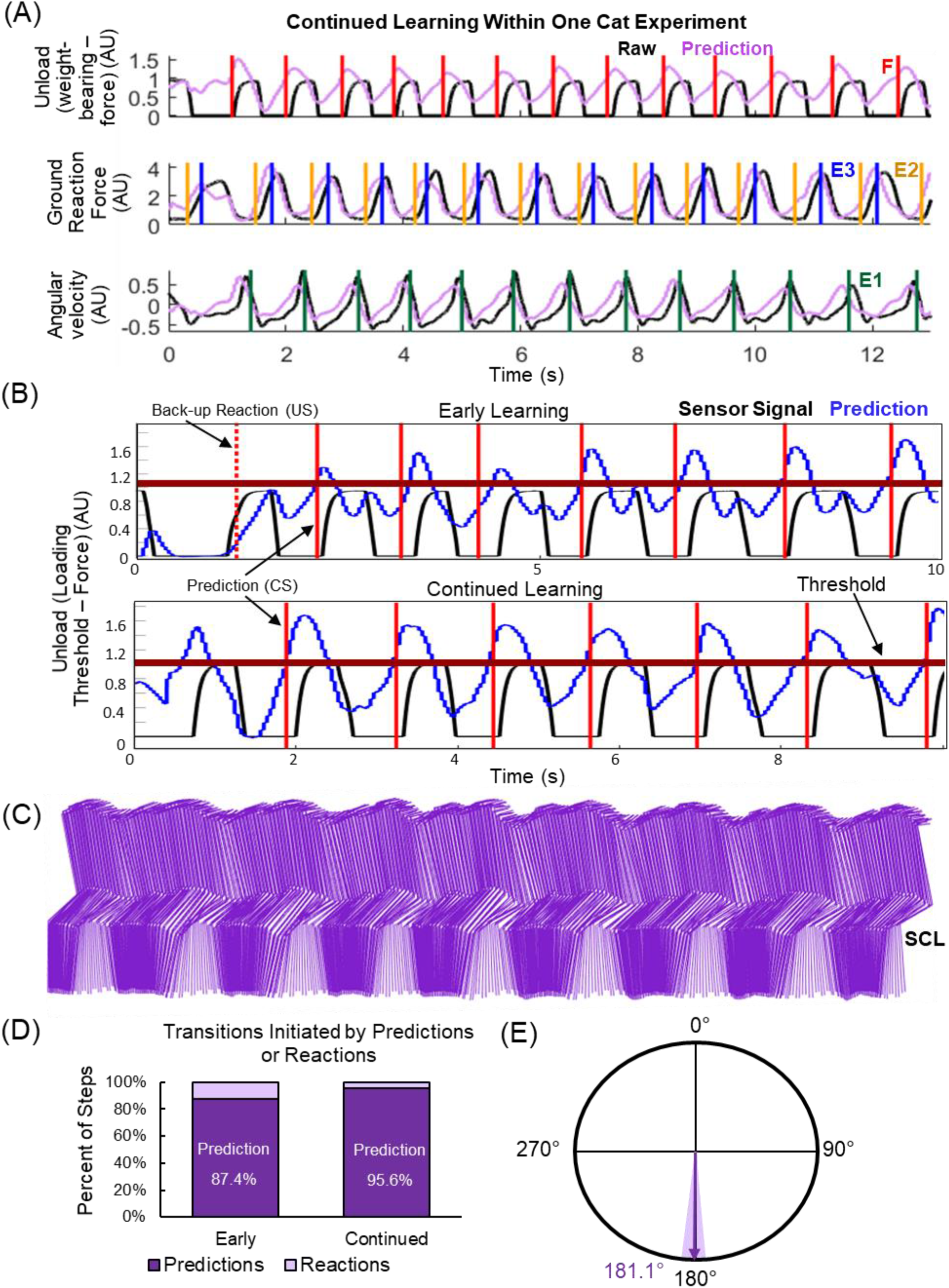
Comparing early learning with later learning. **(A)** Raw sensor signals and learned predictions of unloading, ground reaction force, and angular velocity of the PML (person-moved limb) when learning continued between trials within a single cat experiment. The trigger times for the phases of the walking cycle are marked on their corresponding signal. **(B)** Representative examples of raw and predicted values of the unloading signal during early learning (initialized at 0) and continued learning (after several walking trials). Transitions for the F (early swing) phase made by back-up reactions (dashed vertical) and predictions (solid vertical) are marked. The solid horizontal line indicates the threshold value for predictions of unloading for Pavlovian control. **(C)** Movements produced by the SCL (stimulation-controlled limb) during a trial where learning continued within a single cat experiment. **(D)** Proportion of transitions initiated by predictions and reactions for early learning and continued learning walking trials. **(E)** Average (arrow) and standard deviation (shaded) alternation for trials where learning continued within a cat experiment.

#### Learning continued to initiate prediction-based transitions across several cats and people to produce over-ground walking

The ability of the Pavlovian control to adapt to sudden changes in walking pattern was examined by evaluating the walking trials at the transition between different naïve experimenters walking the PML. As different naïve experimenters took turns to move the PML through the walking cycle, learning quickly acclimated to the new person and their style of walking. Of the 84 transition points between people, 64 did not require a back-up reaction to transition the SCL through the phases of the walking cycle (Figure 7A). Only 5 trials required more than 1 step to adjust to the new naïve experimenter walking the PML before only prediction-triggered transitions occurred. This demonstrated impressively fast adaptations to new environments, resulting in the first personalized and automatically predictive control strategy for a neural prosthesis. The next point of interest was to determine if the settings learned in for Pavlovian control in one animal could transfer and adapt to the next animal. This is analogous to having a new user of a clinical system have their initial settings mirror those of previous users instead of fine tuning the settings from scratch for the new user. Very interestingly, the learned predictions from previous experiments translated well to new cat experiments. In the first walking trial in the new animal, 83.3% of the steps taken did not require a back-up reaction for phase transitions, and 10.0% of steps requiring a back-up reaction were the first step in the trial. The steps in these walking trials were alternating, with an average phase difference of 179.1°±3.2°, which was significantly different from 180° but with a small Cohen’s d effect size (p = 0.026; df = 60; one-sample t-test; Cohen’s d = 0.29; Figure 7B).

**Figure 7.**
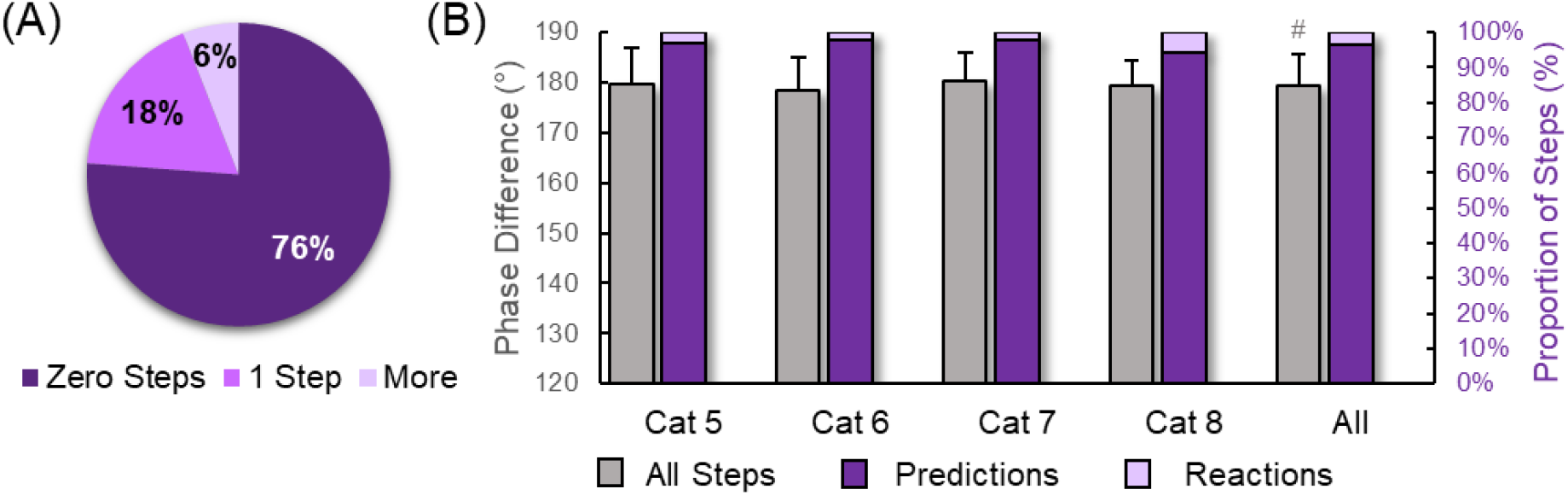
Learning adapts to different subjects during walking. **(A)** Proportion of steps with a back-up reaction following a transition to a new person walking the PML (person-moved limb; n = 84 transitions). **(B)** Alternation and proportion of transitions initiated by predictions and reactions when learning carried over between different cat experiments (n = 61 trials). ^#^p = 0.026.

#### Learning continued to improve across several cats and people to produce over-ground walking

Long-term learning during walking was possible by continuing the learning over several cat experiments with different naïve experimenters taking turns to walk the PML. This provided an excellent representation of day-to-day changes that may occur in the walking patterns produced by the users. The learned predictions triggered phase transitions in more than 91% of the steps taken for all naïve experimenters walking the PML, which was significantly higher than the proportion of prediction initiated transitions in early learning trials (*p* < 0.0001; **Χ**^2^ test Figure 8A). Up to 98.7% of steps were transitioned using predictions (Person B; Figure 8B). On average, these continuing walking trials had a phase difference of 180.8° ± 5.5° (p = 0.113; df = 114; one-sample t-test; Figure 8C). Importantly, there were no missing phase transitions for walking in any trials where learning continued beyond the first learning trial.

**Figure 8.**
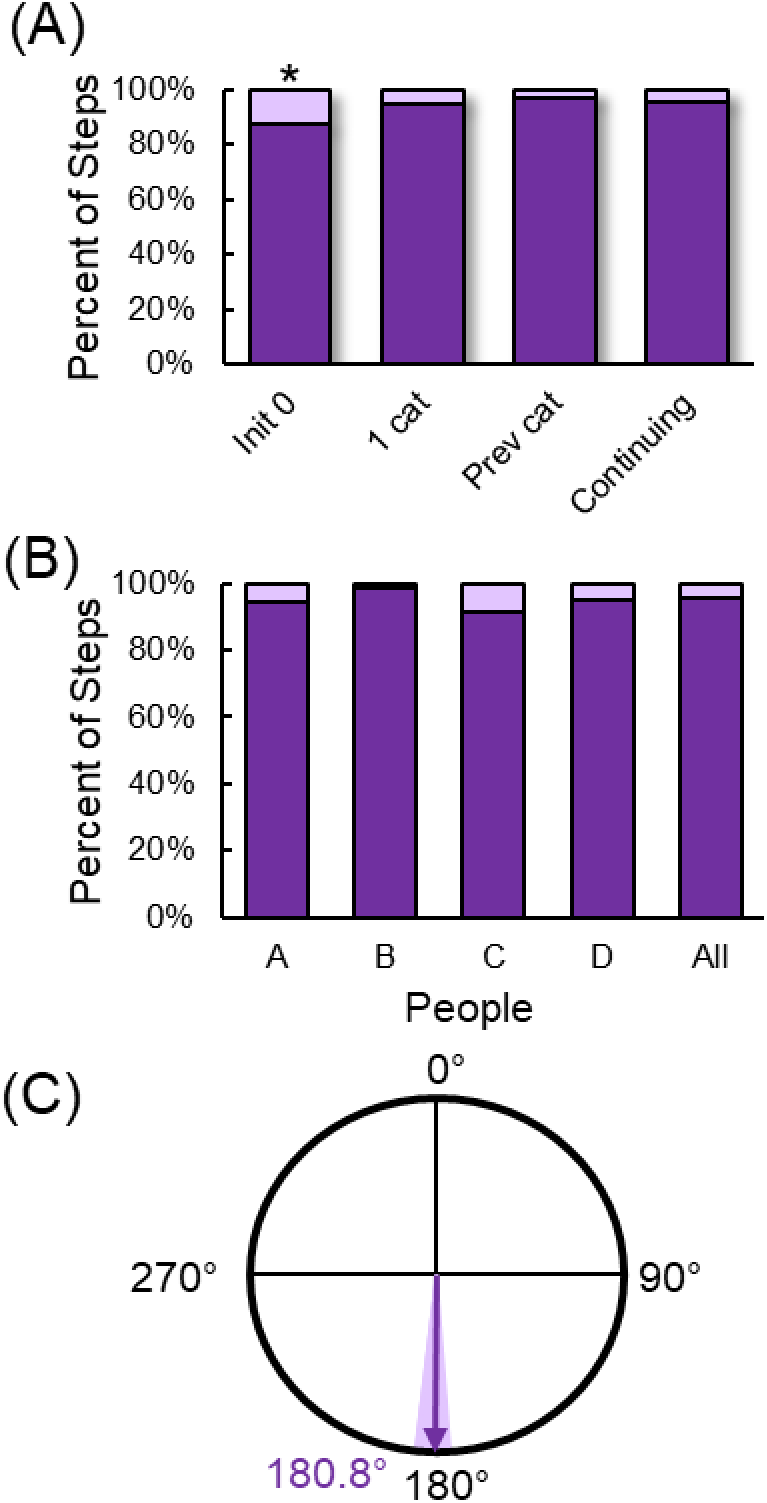
**(A)** Proportion of steps with phase transitions initiated by the learned predictions crossing the threshold or by the back-up reaction at various stages of learning. *p < 0.0001. **(B)** Proportion of steps with phase transitions initiated by the learned predictions or back-up reactions crossing the thresholds for all people walking the PML when learning continued across 5 cat experiments (n = 115 trials). **(D)** Average (arrow) and standard deviation (shaded) of the alternation phase difference of the hind-limbs when walking continued across 5 cat experiments and 4 people walking the PML.

### Pavlovian control recovered from mistakes

Finally, we tested how the learning recovered from perturbations during walking. This is important because the end users of a neural prosthesis may have instances of instability. Different types of intentional mistakes were made by the naïve experimenters walking the PML throughout various stages of learning. The predictions exhibited adaptation to the new and unexpected values of the sensor signals when walking interrupted. Following a mistake, 94.4% (51/54) of the steps that followed had phase transitions triggered by the predicted sensor values (Figure 9). Therefore, not only was the Pavlovian controller able to accommodate multiple users (i.e., cats) and multiple patterns of walking (i.e., different people walking the PML), but it also was able to recover from mistakes made during walking.

**Figure 9.**
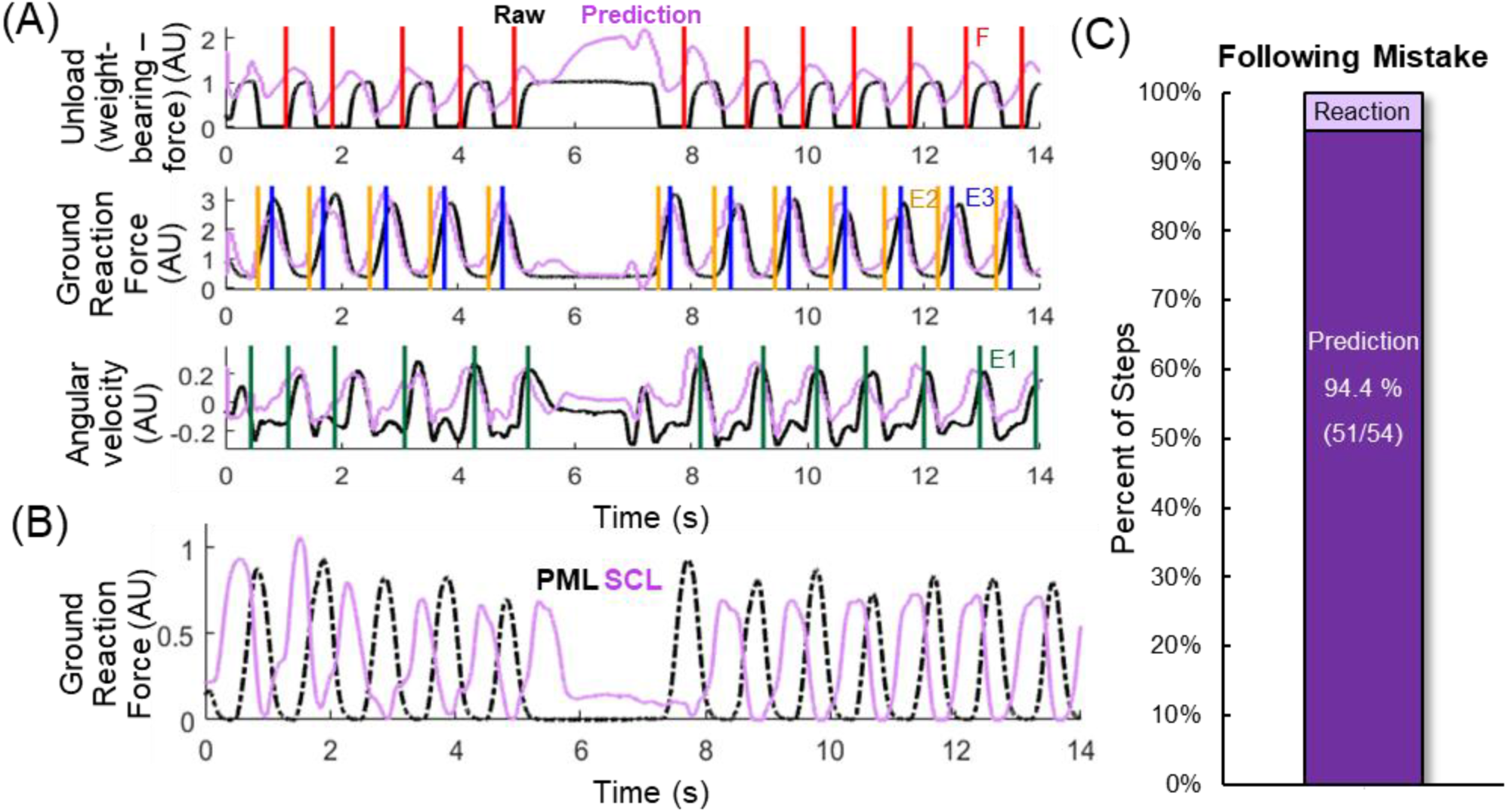
Example of an intentional mistake. **(A)** Raw signals for unloading, ground reaction force, and angular velocity of the PML (person-moved limb) and their corresponding learned predictions. Prediction-initiated transitions are indicated by solid vertical lines. The mistake begins at approximately 5 s and ends near 7.5 s. **(B)** Ground reaction forces produced by the PML (dashed trace) and SCL (stimulation-controlled limb; solid trace). **(C)** Proportion of steps with phase transitions initiated by the learned predictions crossing the threshold or by the back-up reaction following a mistake (n = 54 steps).

## DISCUSSION

The goal of this study was to produce, for the first time, predictive, versatile, alternating, over-ground walking in a model of hemisection SCI using ISMS. The control strategy took advantage of “residual function” and restore over-ground walking in anaesthetized cats. Reinforcement learning was used to learn predictions of walking-relevant sensor values. Pavlovian control used the predicted sensor values and threshold crossings on these predictions to control ISMS such that the “affected limb” is moved to the opposite phase of the walking cycle as the “unaffected limb” in the walking cycle. Pavlovian control can be used across different people walking the “unaffected limb” and throughout different cat experiments without requiring adjustments to the threshold settings. Learning occurred very quickly and consistently produced prediction-driven transitions between the phases of the gait cycle. The learned predictions were also resilient enough to recover quickly following a mistake during walking. Personalized walking was possible for the first time because reinforcement learning acclimated to different people moving the “unaffected limb” and different and cats. This comes in contrast to other approaches where the pattern of walking by the user is dictated by the control algorithm.

### Learning Methods

This study used TOTD to learn GVFs for three cumulants during walking that were used for Pavlovian control. When a GVF crossed a pre-defined threshold, a stimulation response was delivered to move the SCL to the opposite phase of the walking cycle as the PML. The selective Kanerva function approximation method, predictions using GVFs, learning through TOTD, and Pavlovian control are relatively recent advancements made in the field of computing science (Travnik and Pilarski 2017; Sutton et al. 2011; van Seijen et al. 2015; Modayil and Sutton 2014). Selective Kanerva coding was chosen as it has proven to perform well online with a large number of sensors (Travnik and Pilarski 2017). It is also simple to implement and conceptualize. GVFs have proven to be a valuable tool in RL. GVFs allow the prediction of arbitrary signals, which makes RL more powerful and applicable to more problems. In the field of rehabilitation, TD(λ) has been used to produce GVFs for upper-limb prostheses (Pilarski et al. 2012; Pilarski et al. 2013a; 2013b; Sherstan and Pilarski 2014; Edwards et al. 2016). TOTD offers an equivalence to the theoretical forward view of TD learning with negligible increase in computational cost (van Seijen et al. 2015) and has been used to predict the shoulder angle of an upper-limb prosthesis (Travnik and Pilarski 2017). Pavlovian control has successively been used to control switching events of an upper-limb prosthesis in able-bodied study participants (Edwards et al. 2013) and participants with an amputations (Edwards et al. 2016). It has also been used to control the turning off and spinning of a mobile robot (Modayil and Sutton 2014).

Pavlovian control is an appropriate approach to restoring walking in a SCI model because learning the GVFs can occur very rapidly. Since the control strategy only requires the prediction to cross a threshold, online control can be initiated quickly. The learned predictions do not fluctuate nor are largely affected by sudden changes in the raw data, making them more reliable for placing thresholds on for control than the raw signals. Additionally, Pavlovian control does not require exploration of the state space, which is necessary in traditional RL control methods. This is beneficial during walking because exploration of the state space could pose a danger to the user. For example, exploration may produce unsafe movement combinations such as double limb unloading. The state space could be restricted to avoid these dangerous situations, but this would limit the capacity of RL and negate its usefulness. Therefore, Pavlovian control, which uses predictions to drive a fixed stimulation response, is suitable for a repetitive task such as walking.

Pavlovian control also allows for the knowledge of the expert designer to be incorporated into the rules that define the uses of the predictions and the output. This study, for the first time, combined all of these methods and used them to control a neural interface to produce over-ground walking *in vivo*.

### Biological Parallels

Making predictions during a functional task is very useful and is commonly done naturally. For example, during walking, the central nervous system is continuously integrating sensory input from cutaneous receptors on the feet, stretch and loading sensors in the muscles and tendons, as well as visual and vestibular information to maneuver through the environment effectively and safely (Zehr et al. 1997; Zehr and Stein 1999; Donelan and Pearson 2004; Marigold 2008; Mathews et al. 2017). These sensory streams can be used to form short-term predictions that can be used in turn to adapt the gait pattern. Unexpected sensory stimuli result in reflexive changes, and with repetition, adaptation to the sensory stimuli occurs. For example, if an obstacle is placed in front of a cat’s hind-limb during the swing phase causing activation of cutaneous receptors on the dorsum of the paw, the knee will flex further to clear the obstacle (McVea and Pearson 2007). This is a reflexive, or automatic response to the sensory stimulus, which is mediated by the spinal cord. If the obstacle is present for 20 stimuli, the foot will lift higher during swing in anticipation of the obstacle. These effects last over 24 hours in some cases. This long-term adaptation of the gait pattern may be mediated by the cerebellum (Xu et al. 2006). Although this is not exactly an example of Pavlovian control, it demonstrates the usefulness of predictions and how they can be utilized by the nervous system.

Pavlovian control is modelled after classical conditioning. An earlier example described Pavlov’s experiments in dogs where the dogs would salivate when a bell is rung because the ringing became associated with the presentation of food (Pavlov 1883). Another example of classical conditioning is the eye-blink reflex, which has been characterized extensively in rabbits (Kehoe and Macrae 2002; Lepora et al. 2007). In response to a noxious stimulus, such as a puff of air (US), the eye blinks (R). If the puff of air is preceded by a tone (CS), the rabbit blinks just prior to the arrival of the air, protecting the eye. This work is somewhat different from these examples of Pavlovian control because natural movements of one limb do not always dictate the movements of the other limb. However, this concept has similarities to the half-center concept from central pattern generators (Brown 1914). The half-center model of the central pattern generator proposed that the left and right limbs mutually inhibit each other such that when one limb is in flexion, the other must be in extension, and vice versa. The current work incorporated concepts from classical conditioning by also utilizing the sensor information for back-up reactions in the event that the prediction did not reach the threshold in time.

### Relation to Other Control Strategies

Once the thresholds for Pavlovian control were modified after the initial cat experiments, they did not require further modification. Pavlovian control performed significantly better than reaction-based control, providing fewer missed steps and requiring no tuning between transitions different people walking the PML or different cats.

Both the Pavlovian and reaction-based controllers were finite state controllers, which is a concept that has been used previously to produce walking in models of SCI. Finite state control has the advantage of incorporating expert knowledge in a straight-forward manner to define the rules for walking (Popović 1993; Sweeney et al. 2000). Finite state control of surface (Andrews et al. 1988) and intramuscular (Guevremont et al. 2007) FES of the leg muscles used information from ground reaction forces and hip angle to control the transition between the phases of the gait cycle. Previous controllers for ISMS in a model of complete SCI used ground reaction forces and hip angle (Saigal et al. 2004; Holinski et al. 2011; Holinski et al. 2016) or recordings from the dorsal root ganglia (Holinski et al. 2013) to transition the hind-limbs through the different phases (Dalrymple and Mushahwar 2017). Control of epidural stimulation of the spinal cord also utilized electroencephalography recordings from the motor cortex to deliver regional stimulation to the spinal cord to assist with flexion and extension movements in hemisected monkeys (Capogrosso et al. 2016) and people with incomplete SCI (Wagner et al. 2018).

The current study demonstrated that predictions can be learned to initiate transitions between the phases of the gait cycle using only two sensor signals: ground reaction force and angular velocity. These sensors can easily be integrated into a wearable system, as gyroscopes are small microchips and force sensitive resistors can be placed in the soles of shoes (Kirkwood et al. 1989; Kostov et al. 1992). Recent work has demonstrated that kinematic data can be used to identify the phases of the gait cycle during walking (Drnach et al. 2018). They used switched linear dynamical systems (SLDS) to model the joint angle kinematics in healthy people walking on a treadmill. The offline SLDS models were able to label the correct phase of the gait cycle with 84% precision. Future work may incorporate more portable sensors such as goniometers along with online models to build predictions of gait phases.

### Experimental Limitations

The model of a hemisection SCI used in this study enabled thorough testing of the control strategies while avoiding the need for inducing SCIs. It allowed testing of the ability of the control strategies to augment residual function in a controlled manner. This necessitated voluntary control of one hind-limb to be mimicked by a person moving the limb through the walking cycle. This was the first testing of these control strategies, and the outcomes served as a proof-of-concept implementation. Further work may test these control strategies in chronically injured cats, either decerebrate or awake.

A hemisection SCI has more stereotypic functional deficits compared to other injuries such as bilateral contusion SCIs. Although these SCIs are rare, e.g., Brown-Sequard syndrome (Roth et al. 1991; Wirz et al. 2010), the control strategies may be extended to hemiplegia in general, which includes stroke and traumatic brain injury.

The thresholds for Pavlovian control were finalized after initial testing in early experiments. They were chosen based on testing on previously collected data from treadmill stepping (Dalrymple et al 2018) and bench testing on the walkway without a cat. Moderate performance of walking was achieved; however, transitions were improved with changes to the thresholds and the signals on which the thresholds were placed. It is important to note that the learning parameters of the predictions were never changed as they were consistently accurate. Additionally, once the new thresholds were set, they were never again modified. This demonstrates that the initial design decision of where to place the thresholds was important, but once it was finalized no further changes were necessary. The thresholds for Pavlovian control did not require tuning for different people and cats, because the learned predictions acclimated to the changes. However, it may be beneficial to introduce adaptive thresholds in the future, especially if these strategies were to be employed in more variable injury models. Furthermore, the stimulation amplitudes and channels that produced the functional responses remained constant during a walking trial. Future work may introduce a learning strategy that aims to optimize and adapt the stimulation channels and amplitudes in addition to a strategy that controls the timing.

### Future Considerations

Pavlovian control learned predictions for ground reaction force and angular velocity signals; however, other sensor signals could also be used to provide more information about the environment. For example, muscle activity recorded using EMG, joint angles provided by goniometers, or visual information through cameras or infrared sensors could all be recorded and used to acquire more predictions. The addition of sensors (e.g. EMG, goniometers) could be useful to restore walking after variable injuries or to provide information regarding the walking terrain (visual, infrared) to adapt the control strategy. Additional sensors could also be used to provide stability information such as loss of balance, fatigue, and the reliance on the upper body for support. More control rules could be incorporated to predict and correct these safe situations. Furthermore, the addition of sensors is feasible if a state representation method such as selective Kanerva coding is used, as was the case in this work, because it is not affected by the increase in dimensions that plagues traditional tile coding (Travnik and Pilarski 2017).

Pavlovian control can easily be expanded to neuromodulation systems such as deep brain stimulation for various conditions including Parkinson’s disease and depression, neuroprosthetic systems for restoring function after stroke or traumatic brain injury, and exoskeletons and artificial limbs.

## CONCLUSION

Pavlovian control of walking augmented function in a hemisection SCI model. Using predictions of sensor signals during walking, Pavlovian control was resilient to transitions between people walking the limb, between cat experiments, and recovered from mistakes made during walking. Pavlovian control of ISMS has the potential to enhance ambulation capacity greatly, generating alternating, over-ground walking. Very importantly, we have demonstrated, for the first time, that control strategies using intelligent machine learning approaches such as Pavlovian control can reduce the burden of tuning stimulation parameters for controlling a neuroprosthesis. This increases the ease of translation of innovative neural technologies to clinical settings.

This control strategy can also be extended to other injury models and other interventions such as peripheral FES, lower-limb prostheses, and exoskeletons.

## Acknowledgements

The authors thank Adrian Lopera Valle and Amirali Toossi for their assistance with the experiments and Rod Gramlich for building the instrumented walkway used in this study. This work was funded by the Canadian Institutes of Health Research. AND was supported by a Queen Elizabeth II Graduate Student Scholarship and DAR was supported by the University of Alberta Undergraduate Research Initiative. VKM is a Canada Research Chair (Tier 1) in Functional Restoration.

## Author Contributions

VKM secured funding, conceived, and supervised the study. AND and RS formulated the machine learning approach. AND developed and programmed the control strategies. AND, RS, and VKM designed the experimental protocols. AND manufactured the implant arrays. AND and DAR performed the surgeries. VKM performed the procedures associated with implantation of the intraspinal microstimulation electrodes. AND and DAR collected and analyzed the data. AND wrote the initial version of the manuscript and made the figures; all authors contributed to its editing. VKM approved the final version of the manuscript on behalf of the co-authors.

## References

Abbas, J. J., and R. J. Triolo. 1997. “Experimental Evaluation of an Adaptive Feedforward Controller for Use in Functional Neuromuscular Stimulation Systems.” IEEE Transactions on Rehabilitation Engineering: A Publication of the IEEE Engineering in Medicine and Biology Society 5 (1): 12–22.

Anderson, Kim D. 2004. “Targeting Recovery: Priorities of the Spinal Cord-Injured Population.” Journal of Neurotrauma 21 (10): 1371–83. https://doi.org/10.1089/neu.2004.21.1371.

Andrews, B. J., R. H. Baxendale, R. Barnett, G. F. Phillips, T. Yamazaki, J. P. Paul, and P. A. Freeman. 1988. “Hybrid FES Orthosis Incorporating Closed Loop Control and Sensory Feedback.” Journal of Biomedical Engineering 10 (2): 189–95.

Angeli, Claudia A., Maxwell Boakye, Rebekah A. Morton, Justin Vogt, Kristin Benton, Yangshen Chen, Christie K. Ferreira, and Susan J. Harkema. 2018. “Recovery of Over-Ground Walking after Chronic Motor Complete Spinal Cord Injury.” The New England Journal of Medicine 379 (13): 1244–50. https://doi.org/10.1056/NEJMoa1803588.

Bamford, J., R. Lebel, K. Parseyan, and V. Mushahwar. 2016. “The Fabrication, Implantation and Stability of Intraspinal Microwire Arrays in the Spinal Cord of Cat and Rat.” IEEE Transactions on Neural Systems and Rehabilitation Engineering PP (99): 1–1. https://doi.org/10.1109/TNSRE.2016.2555959.

Bhumbra, Gardave S., and Marco Beato. 2018. “Recurrent Excitation between Motoneurones Propagates across Segments and Is Purely Glutamatergic.” PLoS Biology 16 (3): e2003586. https://doi.org/10.1371/journal.pbio.2003586.

Brown, T. G. 1914. “On the Nature of the Fundamental Activity of the Nervous Centres; Together with an Analysis of the Conditioning of Rhythmic Activity in Progression, and a Theory of the Evolution of Function in the Nervous System.” The Journal of Physiology 48 (1): 18–46.

Capogrosso, Marco, Tomislav Milekovic, David Borton, Fabien Wagner, Eduardo Martin Moraud, Jean-Baptiste Mignardot, Nicolas Buse, et al. 2016. “A Brain-Spine Interface Alleviating Gait Deficits after Spinal Cord Injury in Primates.” Nature 539 (7628): 284–88. https://doi.org/10.1038/nature20118.

Carhart, Michael R., Jiping He, Richard Herman, S. D’Luzansky, and Wayne T. Willis. 2004. “Epidural Spinal-Cord Stimulation Facilitates Recovery of Functional Walking Following Incomplete Spinal-Cord Injury.” IEEE Transactions on Neural Systems and Rehabilitation Engineering: A Publication of the IEEE Engineering in Medicine and Biology Society 12 (1): 32–42. https://doi.org/10.1109/TNSRE.2003.822763.

Chang, Sarah R., Rudi Kobetic, Musa L. Audu, Roger D. Quinn, and Ronald J. Triolo. 2015. “Powered Lower-Limb Exoskeletons to Restore Gait for Individuals with Paraplegia – a Review.” Case Orthopaedic Journal 12 (1): 75–80.

Chaplin, E. 1996. “Functional Neuromuscular Stimulation for Mobility in People with Spinal Cord Injuries. The Parastep I System.” The Journal of Spinal Cord Medicine 19 (2): 99–105.

Dalrymple, Ashley N., Dirk G. Everaert, David S. Hu, and Vivian K. Mushahwar. 2018. “A Speed-Adaptive Intraspinal Microstimulation Controller to Restore Weight-Bearing Stepping in a Spinal Cord Hemisection Model.” Journal of Neural Engineering 15 (5): 056023. https://doi.org/10.1088/1741-2552/aad872.

Dalrymple, Ashley N., and Vivian K. Mushahwar. 2017. “Stimulation of the Spinal Cord for the Control of Walking.” In Neuroprosthetics, Volume 8:811–49. Series on Bioengineering and Biomedical Engineering, Volume 8. World Scientific. https://doi.org/10.1142/9789813207158_0025.

Donelan, J. Maxwell, and Keir G. Pearson. 2004. “Contribution of Sensory Feedback to Ongoing Ankle Extensor Activity during the Stance Phase of Walking.” Canadian Journal of Physiology and Pharmacology 82 (8–9): 589–98. https://doi.org/10.1139/y04-043.

Drnach, Luke, Irfan Essa, and Lena H. Ting. 2018. “Identifying Gait Phases from Joint Kinematics during Walking with Switched Linear Dynamical Systems.” BioRxiv, July, 378380. https://doi.org/10.1101/378380.

Edwards, Ann L., Michael R. Dawson, Jacqueline S. Hebert, Craig Sherstan, Richard S. Sutton, K. Ming Chan, and Patrick M. Pilarski. 2016. “Application of Real-Time Machine Learning to Myoelectric Prosthesis Control: A Case Series in Adaptive Switching.” Prosthetics and Orthotics International 40 (5): 573–81. https://doi.org/10.1177/0309364615605373.

Edwards, Ann L., Alexandra Kearney, Michael Rory Dawson, Richard S. Sutton, and Patrick M. Pilarski. 2013. “Temporal-Difference Learning to Assist Human Decision Making during the Control of an Artificial Limb.” ArXiv:1309.4714 [Cs], September. http://arxiv.org/abs/1309.4714.

Ekelem, Andrew, and Michael Goldfarb. 2018. “Supplemental Stimulation Improves Swing Phase Kinematics During Exoskeleton Assisted Gait of SCI Subjects With Severe Muscle Spasticity.” Frontiers in Neuroscience 12: 374. https://doi.org/10.3389/fnins.2018.00374.

Engberg, I., and A. Lundberg. 1969. “An Electromyographic Analysis of Muscular Activity in the Hindlimb of the Cat during Unrestrained Locomotion.” Acta Physiologica Scandinavica 75 (4): 614–30. https://doi.org/10.1111/j.1748-1716.1969.tb04415.x.

Fisekovic, N., and D. B. Popovic. 2001. “New Controller for Functional Electrical Stimulation Systems.” Medical Engineering & Physics 23 (6): 391–99.

Gill, Megan L., Peter J. Grahn, Jonathan S. Calvert, Margaux B. Linde, Igor A. Lavrov, Jeffrey A. Strommen, Lisa A. Beck, et al. 2018. “Neuromodulation of Lumbosacral Spinal Networks Enables Independent Stepping after Complete Paraplegia.” Nature Medicine, September. https://doi.org/10.1038/s41591-018-0175-7.

Goslow, G. E., R. M. Reinking, and D. G. Stuart. 1973. “The Cat Step Cycle: Hind Limb Joint Angles and Muscle Lengths during Unrestrained Locomotion.” Journal of Morphology 141 (1): 1–41. https://doi.org/10.1002/jmor.1051410102.

Guevremont, Lisa, Jonathan A. Norton, and Vivian K. Mushahwar. 2007. “Physiologically Based Controller for Generating Overground Locomotion Using Functional Electrical Stimulation.” Journal of Neurophysiology 97 (3): 2499–2510. https://doi.org/10.1152/jn.01177.2006.

Hardin, Elizabeth, Rudi Kobetic, Lori Murray, Michelle Corado-Ahmed, Gilles Pinault, Jonathan Sakai, Stephanie Nogan Bailey, Chester Ho, and Ronald J. Triolo. 2007. “Walking after Incomplete Spinal Cord Injury Using an Implanted FES System: A Case Report.” Journal of Rehabilitation Research and Development 44 (3): 333–46.

Hofstoetter, Ursula S., Matthias Krenn, Simon M. Danner, Christian Hofer, Helmut Kern, William B. McKay, Winfried Mayr, and Karen Minassian. 2015. “Augmentation of Voluntary Locomotor Activity by Transcutaneous Spinal Cord Stimulation in Motor-Incomplete Spinal Cord-Injured Individuals.” Artificial Organs 39 (10): E176–186. https://doi.org/10.1111/aor.12615.

Holinski, B. J., D. G. Everaert, V. K. Mushahwar, and R. B. Stein. 2013. “Real-Time Control of Walking Using Recordings from Dorsal Root Ganglia.” Journal of Neural Engineering 10 (5): 056008. https://doi.org/10.1088/1741-2560/10/5/056008.

Holinski, B. J., K. A. Mazurek, D. G. Everaert, A. Toossi, A. M. Lucas-Osma, P. Troyk, R. Etienne-Cummings, R. B. Stein, and V. K. Mushahwar. 2016. “Intraspinal Microstimulation Produces Over-Ground Walking in Anesthetized Cats.” Journal of Neural Engineering 13 (5): 056016. https://doi.org/10.1088/1741-2560/13/5/056016.

Holinski, Bradley J., Kevin A. Mazurek, Dirk G. Everaert, Richard B. Stein, and Vivian K. Mushahwar. 2011. “Restoring Stepping after Spinal Cord Injury Using Intraspinal Microstimulation and Novel Control Strategies.” Conference Proceedings: … Annual International Conference of the IEEE Engineering in Medicine and Biology Society. IEEE Engineering in Medicine and Biology Society. Annual Conference 2011: 5798–5801. https://doi.org/10.1109/IEMBS.2011.6091435.

Hunter, J. P., and P. Ashby. 1994. “Segmental Effects of Epidural Spinal Cord Stimulation in Humans.” The Journal of Physiology 474 (3): 407–19.

Johnston, T. E., R. R. Betz, B. T. Smith, B. J. Benda, M. J. Mulcahey, R. Davis, T. P. Houdayer, M. A. Pontari, A. Barriskill, and G. H. Creasey. 2005. “Implantable FES System for Upright Mobility and Bladder and Bowel Function for Individuals with Spinal Cord Injury.” Spinal Cord 43 (12): 713–23. https://doi.org/10.1038/sj.sc.3101797.

Kehoe, E. James, and Michaela Macrae. 2002. “Fundamental Behavioral Methods and Findings in Classical Conditioning.” In A Neuroscientist’s Guide to Classical Conditioning, edited by John W. Moore, 171–231. New York, NY: Springer New York. https://doi.org/10.1007/978-1-4419-8558-3_6.

Kirkwood, C. A., and B. J. Andrews. 1989. “Finite State Control of FES Systems: Application of AI Inductive Learning Techniques.” In Images of the Twenty-First Century. Proceedings of the Annual International Engineering in Medicine and Biology Society, 1020–21 vol.3. https://doi.org/10.1109/IEMBS.1989.96065.

Kirkwood, C. A., B. J. Andrews, and P. Mowforth. 1989. “Automatic Detection of Gait Events: A Case Study Using Inductive Learning Techniques.” Journal of Biomedical Engineering 11 (6): 511–16.

Kobetic, R., R. J. Triolo, and E. B. Marsolais. 1997. “Muscle Selection and Walking Performance of Multichannel FES Systems for Ambulation in Paraplegia.” IEEE Transactions on Rehabilitation Engineering: A Publication of the IEEE Engineering in Medicine and Biology Society 5 (1): 23–29.

Kostov, A., B. J. Andrews, D. B. Popović, R. B. Stein, and W. W. Armstrong. 1995. “Machine Learning in Control of Functional Electrical Stimulation Systems for Locomotion.” IEEE Transactions on Bio-Medical Engineering 42 (6): 541–51. https://doi.org/10.1109/10.387193.

Kostov, A., R. B. Stein, W. W. Armstrong, and M. Thomas. 1992. “Evaluation of Adaptive Logic Networks for Control of Walking in Paralyzed Patients.” In 1992 14th Annual International Conference of the IEEE Engineering in Medicine and Biology Society, 4:1332–34. https://doi.org/10.1109/IEMBS.1992.5761816.

Kunam, Vamsi K., Vinodkumar Velayudhan, Zeshan A. Chaudhry, Matthew Bobinski, Wendy R. K. Smoker, and Deborah L. Reede. 2018. “Incomplete Cord Syndromes: Clinical and Imaging Review.” Radiographics: A Review Publication of the Radiological Society of North America, Inc 38 (4): 1201–22. https://doi.org/10.1148/rg.2018170178.

Lam, Tania, Katherine Pauhl, Amanda Ferguson, Raza N. Malik, BKin, Andrei Krassioukov, and Janice J. Eng. 2015. “Training with Robot-Applied Resistance in People with Motor-Incomplete Spinal Cord Injury: Pilot Study.” Journal of Rehabilitation Research and Development 52 (1): 113–29. https://doi.org/10.1682/JRRD.2014.03.0090.

Lau, Bernice, Lisa Guevremont, and Vivian K. Mushahwar. 2007. “Strategies for Generating Prolonged Functional Standing Using Intramuscular Stimulation or Intraspinal Microstimulation.” IEEE Transactions on Neural Systems and Rehabilitation Engineering: A Publication of the IEEE Engineering in Medicine and Biology Society 15 (2): 273–85. https://doi.org/10.1109/TNSRE.2007.897030.

Lepora, N. F., E. Mavritsaki, J. Porrill, C. H. Yeo, C. Evinger, and P. Dean. 2007. “Evidence from Retractor Bulbi EMG for Linearized Motor Control of Conditioned Nictitating Membrane Responses.” Journal of Neurophysiology 98 (4): 2074–88. https://doi.org/10.1152/jn.00210.2007.

Marigold, Daniel S. 2008. “Role of Peripheral Visual Cues in Online Visual Guidance of Locomotion.” Exercise and Sport Sciences Reviews 36 (3): 145–51. https://doi.org/10.1097/JES.0b013e31817bff72.

Mathews, Miranda A., Aaron J. Camp, and Andrew J. Murray. 2017. “Reviewing the Role of the Efferent Vestibular System in Motor and Vestibular Circuits.” Frontiers in Physiology 8: 552. https://doi.org/10.3389/fphys.2017.00552.

McVea, D. A., and K. G. Pearson. 2007. “Long-Lasting, Context-Dependent Modification of Stepping in the Cat after Repeated Stumbling-Corrective Responses.” Journal of Neurophysiology 97 (1): 659–69. https://doi.org/10.1152/jn.00921.2006.

Modayil, Joseph, and Richard S. Sutton. 2014. “Prediction Driven Behavior: Learning Predictions That Drive Fixed Responses.” 2014. https://www.aaai.org/ocs/index.php/WS/AAAIW14/paper/view/8740.

Moritz, Chet T., Steve I. Perlmutter, and Eberhard E. Fetz. 2008. “Direct Control of Paralysed Muscles by Cortical Neurons.” Nature 456 (7222): 639–42. https://doi.org/10.1038/nature07418.

Morrison, Sarah A., Douglas Lorenz, Carol P. Eskay, Gail F. Forrest, and D. Michele Basso. 2018. “Longitudinal Recovery and Reduced Costs After 120 Sessions of Locomotor Training for Motor Incomplete Spinal Cord Injury.” Archives of Physical Medicine and Rehabilitation 99 (3): 555–62. https://doi.org/10.1016/j.apmr.2017.10.003.

Mushahwar, V. K., D. F. Collins, and A. Prochazka. 2000. “Spinal Cord Microstimulation Generates Functional Limb Movements in Chronically Implanted Cats.” Experimental Neurology 163 (2): 422–29. https://doi.org/10.1006/exnr.2000.7381.

Mushahwar, V. K., and K. W. Horch. 1998. “Selective Activation and Graded Recruitment of Functional Muscle Groups through Spinal Cord Stimulation.” Annals of the New York Academy of Sciences 860 (November): 531–35.

Mushahwar, V. K., and K. W. Horch. 2000. “Selective Activation of Muscle Groups in the Feline Hindlimb through Electrical Microstimulation of the Ventral Lumbo-Sacral Spinal Cord.” IEEE Transactions on Rehabilitation Engineering: A Publication of the IEEE Engineering in Medicine and Biology Society 8 (1): 11–21.

Musselman, Kristin E., Karim Fouad, John E. Misiaszek, and Jaynie F. Yang. 2009. “Training of Walking Skills Overground and on the Treadmill: Case Series on Individuals with Incomplete Spinal Cord Injury.” Physical Therapy 89 (6): 601–11. https://doi.org/10.2522/ptj.20080257.

Pavlov, Ivan Petrovich. 1883. “The Work of the Digestive Glands.” Bristol Medico-Chirurgical Journal (1883) 21 (80). https://www.ncbi.nlm.nih.gov/pmc/articles/PMC5046328/.

Pilarski, P. M., M. R. Dawson, T. Degris, J. P. Carey, and R. S. Sutton. 2012. “Dynamic Switching and Real-Time Machine Learning for Improved Human Control of Assistive Biomedical Robots.” In 2012 4th IEEE RAS EMBS International Conference on Biomedical Robotics and Biomechatronics (BioRob), 296–302. https://doi.org/10.1109/BioRob.2012.6290309.

Pilarski, Patrick M., Michael R. Dawson, Thomas Degris, Jason P. Carey, K. Ming Chan, Jacqueline S. Hebert, and Richard S. Sutton. 2013. “Adaptive Artificial Limbs: A Real-Time Approach to Prediction and Anticipation.” IEEE Robotics Automation Magazine 20 (1): 53–64.

Pilarski, Patrick M., Travis B. Dick, and Richard S. Sutton. 2013. “Real-Time Prediction Learning for the Simultaneous Actuation of Multiple Prosthetic Joints.” IEEE … International Conference on Rehabilitation Robotics: [Proceedings] 2013 (June): 6650435. https://doi.org/10.1109/ICORR.2013.6650435.

Popović, D. B. 1993. “Finite State Model of Locomotion for Functional Electrical Stimulation Systems.” Progress in Brain Research 97: 397–407.

Popović, D., R. B. Stein, N. Oğuztöreli, M. Lebiedowska, and S. Jonić. 1999. “Optimal Control of Walking with Functional Electrical Stimulation: A Computer Simulation Study.” IEEE Transactions on Rehabilitation Engineering: A Publication of the IEEE Engineering in Medicine and Biology Society 7 (1): 69–79.

Qi, H., D. J. Tyler, and D. M. Durand. 1999. “Neurofuzzy Adaptive Controlling of Selective Stimulation for FES: A Case Study.” IEEE Transactions on Rehabilitation Engineering: A Publication of the IEEE Engineering in Medicine and Biology Society 7 (2): 183–92.

Roth, E. J., T. Park, T. Pang, G. M. Yarkony, and M. Y. Lee. 1991. “Traumatic Cervical Brown-Sequard and Brown-Sequard-plus Syndromes: The Spectrum of Presentations and Outcomes.” Paraplegia 29 (9): 582–89. https://doi.org/10.1038/sc.1991.86.

Saigal, R., C. Renzi, and V.K. Mushahwar. 2004. “Intraspinal Microstimulation Generates Functional Movements after Spinal-Cord Injury.” IEEE Transactions on Neural Systems and Rehabilitation Engineering 12 (4): 430–40. https://doi.org/10.1109/TNSRE.2004.837754.

Seijen, Harm H van, and Richard S Sutton. 2014. “True Online TD(Lambda).” In. Vol. 32. Beijing, China.

Seijen, Harm van, A. Rupam Mahmood, Patrick M. Pilarski, Marlos C. Machado, and Richard S. Sutton. 2015. “True Online Temporal-Difference Learning,” December. http://arxiv.org/abs/1512.04087.

Sepulveda, F., M. H. Granat, and A. Cliquet. 1997. “Two Artificial Neural Systems for Generation of Gait Swing by Means of Neuromuscular Electrical Stimulation.” Medical Engineering & Physics 19 (1): 21–28.

Sherstan, Craig, and Patrick Pilarski. 2014. “Multilayer General Value Functinos for Robotic Prediction and Control.” In, 6. Chicago, IL,vUSA.

Skinner, BF. 1963. “Operant Behavior.” American Psychologist 18 (8): 503–15.

“Spinal Cord Injury (SCI) 2017 Facts and Figures at a Glance.” 2017. The Journal of Spinal Cord Medicine 40 (6): 872–73. https://doi.org/10.1080/10790268.2017.1379938.

Staddon, J. E. R., and D. T. Cerutti. 2003. “Operant Conditioning.” Annual Review of Psychology 54: 115–44. https://doi.org/10.1146/annurev.psych.54.101601.145124.

Sutton, Richard S. 1988. “Learning to Predict by the Methods of Temporal Differences.” Machine Learning 3 (1): 9–44. https://doi.org/10.1007/BF00115009.

Sutton, Richard S, and Andrew G Barto. 2018. Reinforcement Learning: An Introduction. 2nd ed. Adaptive Computation and Machine Learning Series. Cambridge, MA: MIT Press.

Sutton, Richard S, Joseph Modayil, Michael Delp, Thomas Degris, Patrick M Pilarski, Adam White, and Doina Precup. 2011. “Horde: A Scalable Real-Time Architecture for Learning Knowledge from Unsupervised Sensorimotor Interaction,” 8.

Sweeney, P. C., G. M. Lyons, and P. H. Veltink. 2000. “Finite State Control of Functional Electrical Stimulation for the Rehabilitation of Gait.” Medical & Biological Engineering & Computing 38 (2): 121–26.

Tong, K. Y., and M. H. Granat. 1999. “Gait Control System for Functional Electrical Stimulation Using Neural Networks.” Medical & Biological Engineering & Computing 37 (1): 35–41.

Travnik, Jaden B., and Patrick M. Pilarski. 2017. “Representing High-Dimensional Data to Intelligent Prostheses and Other Wearable Assistive Robots: A First Comparison of Tile Coding and Selective Kanerva Coding.” IEEE … International Conference on Rehabilitation Robotics: [Proceedings] 2017: 1443–50. https://doi.org/10.1109/ICORR.2017.8009451.

Vanderhorst, V. G., and G. Holstege. 1997. “Organization of Lumbosacral Motoneuronal Cell Groups Innervating Hindlimb, Pelvic Floor, and Axial Muscles in the Cat.” The Journal of Comparative Neurology 382 (1): 46–76.

Wagner, Fabien B., Jean-Baptiste Mignardot, Camille G. Le Goff-Mignardot, Robin Demesmaeker, Salif Komi, Marco Capogrosso, Andreas Rowald, et al. 2018. “Targeted Neurotechnology Restores Walking in Humans with Spinal Cord Injury.” Nature 563 (7729): 65–71. https://doi.org/10.1038/s41586-018-0649-2.

Wen, Yue, Jennie Si, Andrea Brandt, Xiang Gao, and He Huang. 2019. “Online Reinforcement Learning Control for the Personalization of a Robotic Knee Prosthesis.” IEEE Transactions on Cybernetics, January. https://doi.org/10.1109/TCYB.2019.2890974.

White, Adam. 2015. “Developing a Predictive Approach to Knowledge.” Edmonton, AB, Canada: University of Alberta.

Williamson, R., and B. J. Andrews. 2000. “Gait Event Detection for FES Using Accelerometers and Supervised Machine Learning.” IEEE Transactions on Rehabilitation Engineering: A Publication of the IEEE Engineering in Medicine and Biology Society 8 (3): 312–19.

Wirz, M., B. Zörner, R. Rupp, and V. Dietz. 2010. “Outcome after Incomplete Spinal Cord Injury: Central Cord versus Brown-Sequard Syndrome.” Spinal Cord 48 (5): 407–14. https://doi.org/10.1038/sc.2009.149.

Xu, Duo, Tao Liu, James Ashe, and Khalafalla O. Bushara. 2006. “Role of the Olivo-Cerebellar System in Timing.” The Journal of Neuroscience: The Official Journal of the Society for Neuroscience 26 (22): 5990–95. https://doi.org/10.1523/JNEUROSCI.0038-06.2006.

Zehr, E. P., T. Komiyama, and R. B. Stein. 1997. “Cutaneous Reflexes during Human Gait: Electromyographic and Kinematic Responses to Electrical Stimulation.” Journal of Neurophysiology 77 (6): 3311–25. https://doi.org/10.1152/jn.1997.77.6.3311.

Zehr, E. P., and R. B. Stein. 1999. “What Functions Do Reflexes Serve during Human Locomotion?” Progress in Neurobiology 58 (2): 185–205.

